# Coherent cortical representations develop after experience via feedforward-recurrent circuit alignment

**DOI:** 10.1101/2023.07.09.547747

**Authors:** Augusto Abel Lempel, Sigrid Trägenap, Clara Tepohl, Matthias Kaschube, David Fitzpatrick

## Abstract

Sensory cortical areas guide behavior by transforming stimulus-driven inputs into selective responses representing relevant features. A classic example is the representation of edge orientations in the visual cortex^1–4^, where layer 4 (L4) neurons co-activated by an orientation provide feedforward inputs to specific functional modules in layer 2/3 (L2/3) that share strong recurrent connections^5–7^. The aligned state of feedforward-recurrent interactions is critical for amplifying selective cortical responses^8–12^, but how it develops remains unclear. Using simultaneous electrophysiology and calcium imaging in visually naïve animals we find that coactivity of L4 neurons and L2/3 modular responses elicited by oriented gratings lacks the tight relationship to orientation preference found in experienced animals. One factor that could contribute to the lack of functionally specific coactivity is high variability in naïve L4 neuron responses that decreases significantly following experience. But a computational model of feedforward-recurrent interaction suggests that high variability alone is insufficient to explain the naïve state and provides a biological signature of feedforward-recurrent misalignment that we confirm with whole-cell recordings: dynamic changes in orientation tuning of L2/3 subthreshold responses shortly after stimulus onset. In conclusion, we provide diverse evidence for a realignment of feedforward-recurrent interactions following experience that is critical for building reliable sensory representations with interlaminar temporal coherence.

## Main

Sensory inputs drive cortical functional networks to produce well-organized representations of stimulus features critical for behavior. These representations rely on constructive interactions between stimulus-evoked inputs and the cortical circuit to support highly reliable spatiotemporal population activity patterns. An extensively studied example is the spatially organized – modular – representation of orientation in layers 2/3 (L2/3) of the primary visual cortex^1–4^. L2/3 neurons receive orientation-selective feedforward inputs from neurons in layer 4 (L4), which acquire their selectivity through the convergence of thalamic inputs^1,13,14^, and share strong lateral recurrent connections capable of amplifying specific input patterns^12,15,16^. Recurrent connections selectively link L2/3 modules with similar orientation preferences^5–7^, such that L4 inputs evoked by particular orientations align with patterns amplified by recurrent L2/3 circuits and drive highly coherent (correlated) responses across layers. This aligned state of feedforward-recurrent interactions can increase response selectivity and reduce variability, facilitating reliable discrimination across stimuli^8–12^.

While our understanding of the organization of feedforward-recurrent interactions in the mature cortex is far from complete, even less is known about how these interactions are established during development. Their foundation is thought to be laid out early in life by activity-independent^17–19^ and activity-dependent^19,20^ endogenous mechanisms, and this is consistent with the observation that the initial orientation tuning at eye opening shares structural features of the mature representation of edge orientation^21–23^. However, there are also striking immaturities in the visual responses evident at eye opening, including highly unreliable responses for repeated presentations (trials) of the same stimulus^23^, and a mismatch in preferred orientation when stimulating the two eyes separately^21^, suggesting the possibility that feedforward-recurrent interactions may still be relatively immature prior to the onset of visual experience.

We examined the early development of feedforward and recurrent interactions in the visual cortex of the ferret, an animal model with a well-developed modular representation of orientation and a delay in the onset of experience through open eyes following birth that facilitates understanding the role of visual experience in cortical development. Our results emphasize that the onset of visual experience initiates a dynamic feedforward-recurrent alignment process critical for constructing a reliable and spatiotemporally coherent representation of stimulus orientation.

## Results

### Responses to oriented stimuli become coherent across layers after eye opening

To investigate developmental changes in the functional organization of feedforward and recurrent interactions that may contribute to the increased reliability of the representation of orientation, we measured the degree of coactivity of individual neurons in L4 and L2/3 with the modular patterns of network activity in L2/3 in response to oriented gratings in visually naïve and experienced ferrets. To this end, we combined simultaneous trial-to-trial measurements of extracellular single-unit spiking responses in these layers with epi-scale calcium imaging of L2/3 modular responses in a 1 mm radius surrounding the electrode location while presenting oriented, drifting gratings (Fig. 1A). For electrophysiology, layers were identified using sink-source analysis of local field potentials (Fig. S1)^24–26^. Single units for which no stimulus elicited a trial-averaged response above 2 Hz were deemed unresponsive and not included in any analysis. For all responsive units, we then computed a non-parametric correlation between single-unit and modular responses, which we call unit-to-modular coactivity (see Methods; Examples are shown in Figure 1B). Finally, the strength of such coactivity was quantified for each single unit as its mean absolute value across imaging pixels and compared across layers in naïve (post-natal day (P)27-31) and experienced ferrets (older than P36 and at least five days after eye opening; Fig. 1C).

**Figure 1:**
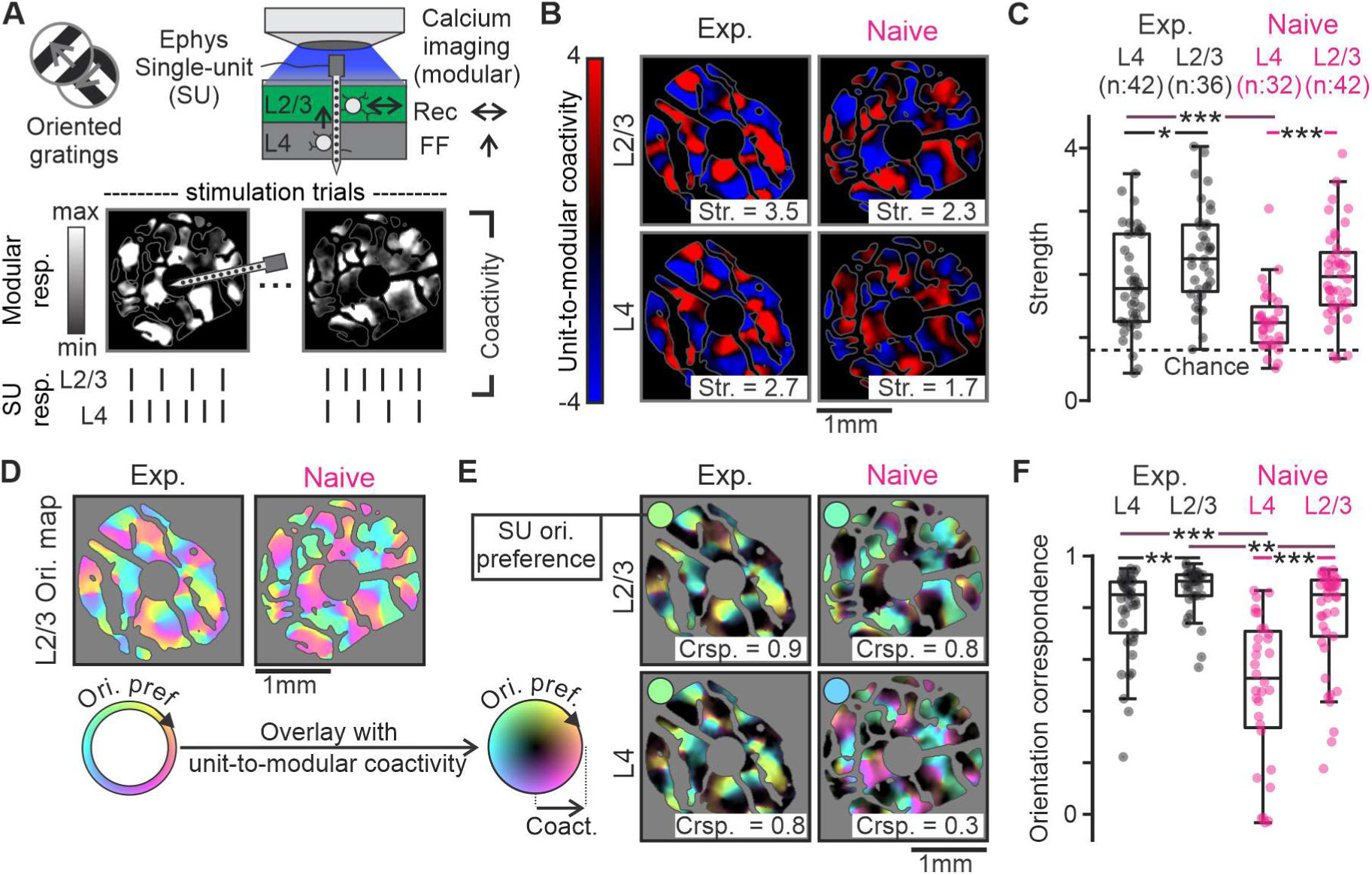
Coactivity of layer 4 (L4) single units with modular layer 2/3 (L2/3) responses undergoes dramatic changes following experience. **A –** Simultaneous epi-scale calcium imaging and extracellular electrophysiology across layers were performed while presenting squared-wave oriented, drifting gratings. Trial-to-trial coactivity between single-unit (SU) spiking and modular L2/3 calcium responses was computed (see Methods). The area immediately surrounding the electrode position (< 250 μm) was not considered. **B –** Example unit-to-modular coactivity for single units recorded in L2/3 or L4 of an experienced and a visually naïve ferret. In our measure of coactivity (see Methods), 1 and -1 indicate that coactivity between SU spiking and pixel calcium responses is 1 standard deviation above and below, respectively, from a shufied (random) distribution. Coactivity strength (Str.) was calculated as the mean absolute coactivity value across pixels in the imaging data. Strength value represents the overall deviation of coactivity, either positive or negative, from shufied distribution, expressed in standard deviations. **C –** Distribution of unit-to-modular coactivity strength for single units recorded in each layer and age group. Dashed line represents chance expectation (∼0.8). *: p < 0.05. ***: p < 0.001. Wilcoxon rank-sum test. **D –** Distribution of L2/3 orientation preferences (orientation map) in the fields of view corresponding to example data shown in B. **E –** Overlay of unit-to-modular coactivity shown in B with the orientation maps shown in D. The preferred orientation of the single units (color circles) is similar to the coactive L2/3 modules’ preferences except for the naïve L4 unit. Orientation correspondence (Crsp.) quantifies these orientation preference similarities (see Methods). **F –** Same as C for orientation correspondence. **: p < 0.01. ***: p < 0.001.

As expected, in experienced animals with highly reliable responses, units in L2/3 show, on average, strong coactivity with modular responses in L2/3 (2.3), and units in L4 show almost equally strong coactivity (1.9. Experienced L4 vs L2/3, p = 0.02). Notably, in naïve animals, which display high trial-to-trial response variability, L2/3 units still exhibit strong coactivity with modular responses with an average value similar to experienced animals (2. Naïve vs Experienced L2/3, p = 0.09). In contrast, L4 units in naïve animals show much weaker coactivity with L2/3 modular responses (1.3) differing significantly from L4 units in experienced animals (Naïve vs Experienced L4, p < 0.001), as well as L2/3 units in the naïve cortex (Naïve L4 vs L2/3, p < 0.001). Note that these laminar differences in coactivity in naïve animals are not due to differences in overall responsiveness (average single-unit response across all grating stimuli: L2/3: 2.47 Hz. L4: 2.71 Hz.).

Our analysis so far demonstrates a lack of maturity in the strength of L4-to-L2/3 (interlaminar) coactivity elicited by oriented gratings before experience and led us to consider the factors that could account for this change. Could this be explained by experience-driven strengthening of stable, preexisting patterns of feedforward-recurrent modular interactions, or are there fundamental changes in the orientation specificity of feedforward-recurrent modular relationships that contribute to the enhanced coactivity?

To explore the possibility of developmental changes in the specificity of feedforward-recurrent modular relationships, we computed the spatial pattern (map) of orientation preference from trial-averaged calcium responses (map examples shown in Figure 1D) and overlay it with the spatial structure of unit-to-modular coactivity (Fig. 1E). For units in the experienced cortex and L2/3 units in naïve animals, coactive modules display orientation preferences similar to that of the unit (Fig. 1E left and upper right). In contrast, for many L4 units in naïve animals, the orientation preferences of coactive modules are not well matched with the L4 unit’s preference (Fig. 1E lower right). To quantify this, we correlated, for each unit, the spatial profile of unit-to-modular coactivity (Fig. 1B) with that of the co-tuning between each pixel and the unit (see methods for details). We call this correlation orientation correspondence (Fig. 1F).

We found that orientation correspondence, while stronger in experienced animals, as expected (average; L4: 0.79; L2/3: 0.87), and strong also for L2/3 units in the naïve cortex (average; 0.76), tended to be much weaker for many naïve L4 single units (average: 0.49. Naïve vs Experienced L4 and Naïve L4 vs L2/3, p < 0.001) (Fig. 1F). Thus, many L4 units in the visually naïve cortex exhibit a much less specific functional coupling (coactivity) with the representation of orientation in L2/3 compared to the experienced cortex. The orientation preference mismatch between naïve L4 units and co-tuned L2/3 modules could not be explained by uncertainty in our estimation of units’ preference due to response variability (Fig. S2). Moreover, naïve L4 units showed lower orientation correspondence than experienced L4 units and naïve L2/3 units with comparable levels of coactivity strength, indicating a degree of independence between these two properties of coactivity (Fig. S3).

Overall, these results point to significant circuit changes that coincide with the emergence of a reliable representation of orientation following visual experience: the coactivity between L4 neurons providing feedforward inputs to L2/3 and modular L2/3 responses becomes stronger and better correlated with orientation preference, indicating increased coherence of responses across layers. This suggests that at the onset of visual experience, the recurrent network in L2/3 has the capacity to amplify poorly correlated signals from L4 to support an initial modular map of orientation. But they also suggest that the immaturities in feedforward-recurrent interactions, as manifested in the strength and orientation correspondence of interlaminar coactivity, could contribute to the weaker tuning and reliability of responses in visually naïve animals.

### The reliability of L4 units’ orientation responses increases following experience

One important factor that could contribute to the immature coactivity of L4 units with the modular patterns of activity in L2/3 at eye opening is the reliability of L4 units’ responses to visual stimulation. To probe this factor, we first computed tuning curves for single-unit responses (Examples shown in Fig. 2A) and quantified orientation discriminability as Cohen’s D between single-trial responses to the preferred and orthogonal orientations (Fig. 2B). The responses of L2/3 and L4 units in visually naïve animals exhibit significantly less discriminability than the corresponding neurons in the experienced animals (L4 naïve = 0.98; L4 experienced = 2.9; L2/3 naïve = 1.73; L2/3 experienced = 3.13. p < 0.001 for both age comparisons). Thus, lack of L4 reliability is likely one contributor to the immature interlaminar coactivity at eye opening.

**Figure 2:**
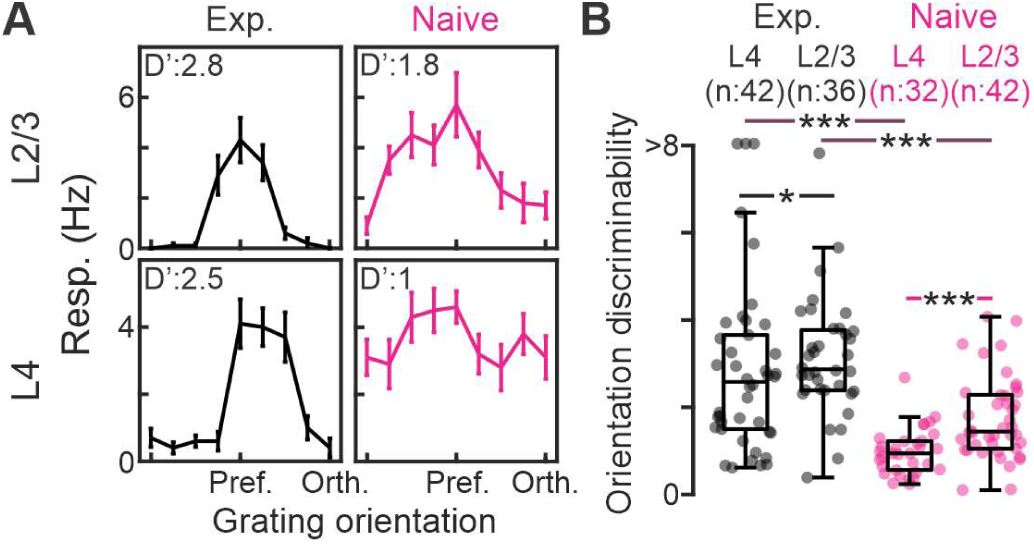
Layer 4 (L4) units orientation discriminability increases dramatically following experience. **A –** Examples of orientation tuning curves of single-unit responses recorded in layer 2/3 (L2/3) and L4 of an experienced and a visually naïve ferret. D’: orientation discriminability. Error bars: standard error of the mean across 8 trials. **B –** Distribution of orientation discriminability across single units recorded in each layer and age group. In visually naïve animals, L4 units exhibit significantly lower orientation discriminability than L2/3 neurons. In experienced animals, discriminability values for both layers become higher and more similar between layers. *: p < 0.05. ***: p < 0.001. Wilcoxon rank-sum test.

Our data also shows enhanced discriminability of L2/3 neurons relative to L4 at eye opening (p < 0.001), in line with previous observations of laminar differences in trial-averaged tuning curves^27^. This, together with the relatively high specificity in the L2/3 unit-to-modular coactivity at eye opening, is consistent with the L2/3 recurrent network amplifying orientation-tuned signals from unreliable L4 inputs at this age. But, does the increase in reliability of L4 responses alone explain the maturation of interlaminar coactivity following experience?

### Feedforward-recurrent alignment can strongly affect cortical response properties

It is conceivable that improvements in the strength and specificity of coactivity between L4 and L2/3 depend not only on L4 response reliability but also on the ability of the L2/3 circuits to amplify their feedforward inputs. Proposed mechanisms for amplifying input signals in recurrent networks are selective^8,23,28–30^ and balanced^31–33^ amplification, both of which are highly input-pattern specific: the strength of amplification depends sensitively on how well an input can constructively interact with the lateral feedback it induces via the recurrent network. The condition for such constructive interaction has been termed feedforward-recurrent alignment, and its experience-dependent improvement has been suggested to play an important role in the development of reliable cortical responses^23^.

To explore a possible role of feedforward-recurrent alignment in addition to enhanced input reliability, we devised a computational firing-rate model of the L2/3 recurrent network and the inputs it receives from L4 (Fig. 3). Its recurrent connectivity (Fig. 3A) was constrained to produce core features of early spontaneous activity when driven by broadly distributed inputs, specifically its modular (Fig. 3B) and low dimensional organization (Fig. S4)^15,23,34^, which is achieved by assuming structured local excitation–lateral inhibition (LE/LI) interactions with mild heterogeneity across cortex^15,16^ (see Methods). Driving each unit in this network with feedforward input of varying discriminability and spatial organization of orientation preference allowed us to systematically study how the network’s response properties depend on the degree of input reliability (Fig. 3C) and feedforward-recurrent alignment (Fig. 3D). For all input conditions, the model network produced modular activity patterns on a trial-to-trial basis (Fig. S5). Analyzing these network responses revealed that their reliability as well as the strength and specificity of their coactivity with feedforward inputs all increase with increases in both input reliability and alignment, with the largest improvements occurring when both factors change (Fig. 3E-G). Moreover, for randomly organized inputs (alignment = 0), not only does the network’s reliability and the input-network coactivity improve very little when increasing input reliability (Fig. 3E, F), but the network also fails to enhance reliability relative to its input (Fig. S6). Finally, low values of orientation correspondence of input-to-network coactivity, as suggested by our data (Fig. 1F), not only require weakly reliable feedforward inputs (Fig. 3G) whose trial-to-trial variability is correlated across units^35–38^ (Fig. S7), but also weak feedforward-recurrent alignment (Fig. 3G). In particular, when input reliability is low and there is very weak alignment (0.1), we observe differences in input-to-network and unit-to-network coactivity orientation correspondence (Fig. S7) similar to that between L4 and L2/3 unit-to-modular coactivity in naïve animals (Fig. 1F). Thus, these results suggest that feedforward-recurrent alignment is poor in the visually naïve cortex and that its improvement can strongly impact the emergence of a robust and specific functional coupling (coherence) between L4 and L2/3 driving highly reliable responses after experience.

**Figure 3:**
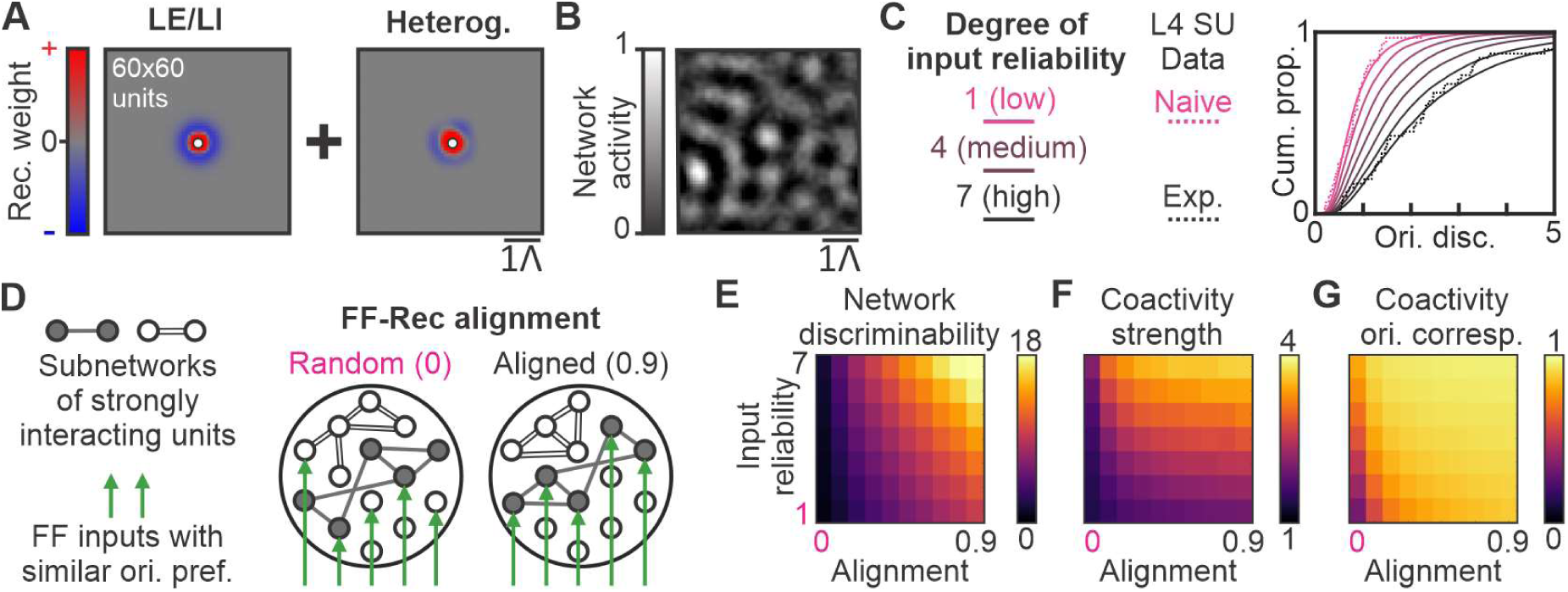
Computational modeling reveals the effects of feedforward-recurrent alignment and input reliability on recurrent network responses. **A –** Recurrent connections were constructed to exhibit a local excitation / lateral inhibition (LE/LI) spatial profile, shown on the left for a unit at the location marked by a white dot. This connectivity was randomly perturbed by adding heterogeneous connections across space (right). Scale bar indicates the characteristic wavelength Λ of activity patterns resulting from such LE/LI connectivity. **B –** Example network activity pattern elicited by spatially weakly smoothed unstructured input. **C –** Definition of input reliability. Pink curves: Cumulative distribution of orientation discriminability of modeled feedforward inputs (solid) matched to the distribution of discriminability of layer 4 (L4) single-unit (SU) data in naïve animals (dotted; same data as in Fig. 2B). Black curves: same, but matched to the distribution in experienced animals. Different solid curves from pink to black show linear interpolations between these two distributions from low (degree 1) to high (degree 7) input reliability. **D –** Feedforward-recurrent alignment. Schematic of the organization of feedforward input with similar orientation preference relative to recurrent connectivity for the random (left) and aligned (right) conditions. **E –** Orientation discriminability of network responses as a function of the degree of input reliability (from 1-7, C) and feedforward-recurrent alignment. **F and G –** Same as E for input-to-network coactivity strength and orientation correspondence.

### Temporal consistency of L2/3 orientation responses increases following experience, as predicted by the development of feedforward-recurrent alignment

Seeing its strong effect on reliable response properties in the model, we next asked whether there is evidence for a developmental improvement of feedforward-recurrent alignment of orientation responses in V1 following the onset of visual experience. To derive a critical prediction from our model, we reasoned that, within the context of single unit orientation tuning, the hallmark of feedforward-recurrent alignment is consistency between the tuning provided by the feedforward inputs and the net tuning of the input received via recurrent connections. If the feedforward drive arrives prior to input fed back by the recurrent network, as previous experiments indicate^39^, and both inputs deviate significantly in their preferred orientation, this could lead to shifts in the unit’s preferred orientation for sustained stimulus presentation. To study this possibility in our model, we analyzed single-unit responses at different latencies following stimulus onset and compared their orientation preference with that of near steady state responses at the end of the simulation period (60τ, Fig. 4A-B). For very weak alignment (0.1), short-latency responses show the largest mismatch in orientation tuning with that at the end of the simulation period (Fig. 4A and S8), indicating that the recurrent inputs exhibit a tuning that is distinct from that of the feedforward inputs. This mismatch is greatly reduced with strong alignment (0.9) (Fig. 4A and S8), as now the orientation preferences of feedforward and recurrent inputs better match each other. Importantly, this mismatch is virtually unaffected by the degree of input reliability (Fig. 4B), indicating the model’s prediction regarding the state of alignment is specific.

**Figure 4:**
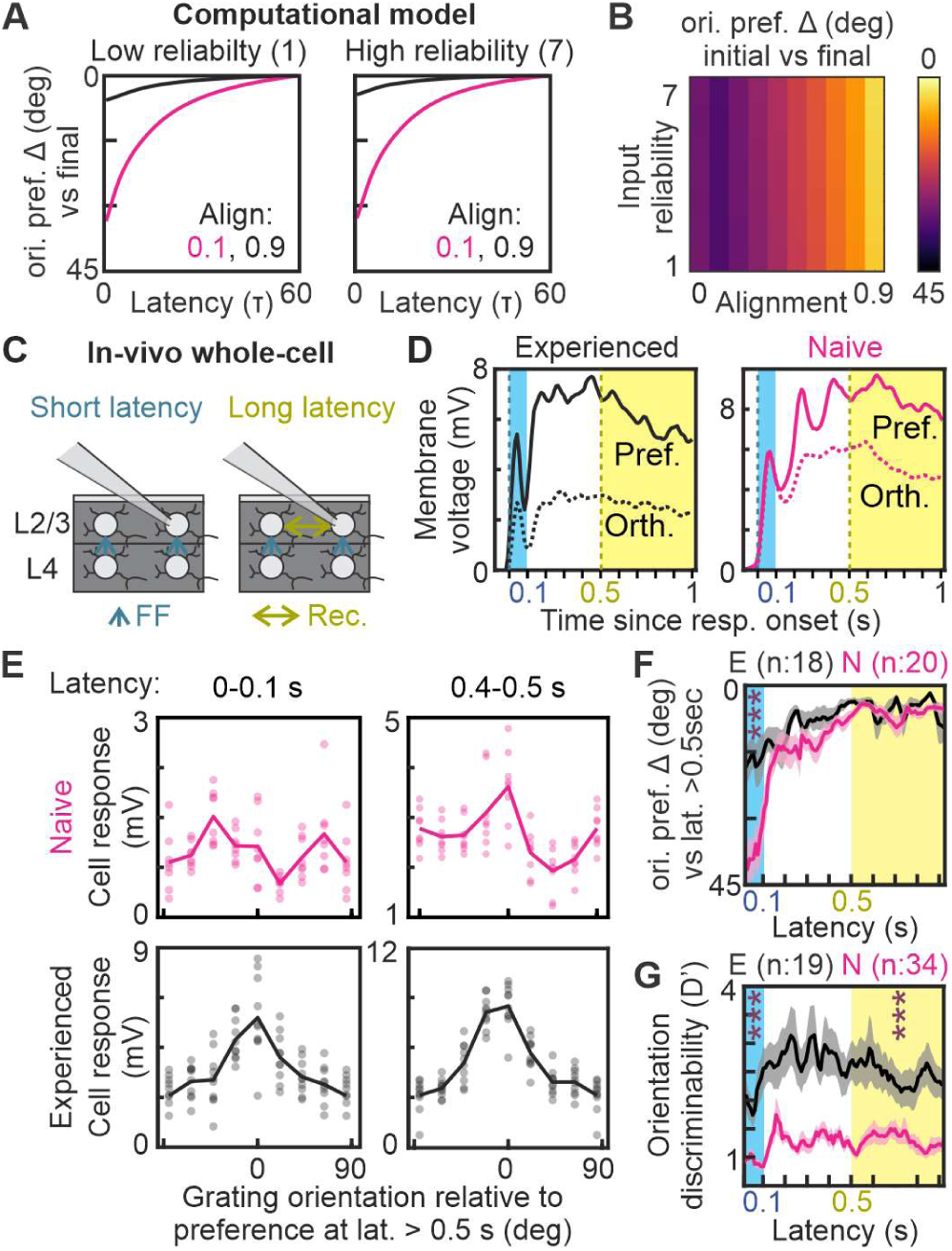
Increased temporal consistency of orientation preference in layer 2/3 (L2/3) following experience confirms predictions of the feedforward-recurrent alignment model. **A –** Line plots of orientation preference mismatch (Δ) between the response at time t following stimulus onset and the final (near steady-state) response (at 60T) for the recurrent network from Fig. 3. The left and right plots show data from model instances with low and high (7) input reliability, respectively. Pink and black lines represent model instances with low (0.1) and high (0.9) feedforward-recurrent alignment, respectively. **B –** Orientation preference mismatch Δ between the initial (t = 1τ) and final (t = 60τ) model response as a function of input reliability and feedforward-recurrent alignment. **C –** In-vivo whole-cell recordings provide data to test the model’s prediction. Short-latency responses (light blue) are expected to reflect the tuning properties of feedforward inputs. The effects of recurrent interactions modulate long-latency, steady-state (yellow) responses. **D-** Example sub-threshold responses to gratings at the preferred (solid) and orthogonal (dotted) orientations as a function of latency, relative to response onset, for L2/3 cells in an experienced and a naïve ferret. **E –** Examples of trial-averaged tuning curves for short and longer-latency sub-threshold responses for L2/3 cells recorded in an experienced and a visually naïve ferret. Line represents trial-averaged tuning curve and dots represent individual trial responses. **F –** Line plots of orientation preference mismatch (Δ), relative to long-latency steady-state responses (latency > 0.5 sec), for L2/3 responses at different latencies in naïve (pink) and experienced (black) ferrets. Shaded region represents standard deviation across cells. ***: p < 0.001 across age. **G –** Same as F for orientation discriminability.

To test these predictions *in vivo*, we performed whole-cell recordings (Fig. 4C) in visually naïve and experienced animals and analyzed the dynamics of sub-threshold responses for different latencies after stimulus onset. For each cell, we computed the onset latency of responses relative to stimulus onset, which was slightly shorter after experience (Mean latency; Naïve: 55 ms. Experienced: 42 ms. p = 0.03). We then computed a “full response” as a temporal average across 1 sec following response onset. In addition, we analyzed responses using a sliding 50 ms temporal average window centered at different latencies from response onset (Fig. 4D).

In the following analysis of orientation tuning dynamics, we focus on cells with reliable orientation preference, i.e. discriminability (D’ > 1), in their full response (20/34 in naïve and 18/19 in experienced animals). To assess the temporal stability of orientation preference, we computed, for each neuron, the orientation tuning curve across latencies (See examples in Figure 4E). We then calculated the orientation preference mismatch between responses at different latencies and that after 500 ms from response onset, when tuning is stable (Fig. 4F). Responses during the first 100 ms following onset display a large orientation preference mismatch in naïve L2/3 neurons that is reduced after experience (p < 0.001), indicating weak levels of feedforward-recurrent alignment for orientation at eye opening that matures following experience. Orientation mismatch and its developmental reduction are not explained by uncertainty in our estimation of orientation preference due to response variability (Fig. S9). Altogether, our modeling and experimental results indicate that poorly aligned feedforward-recurrent interactions drive response dynamics lacking temporal coherence in orientation preference, and this critical property matures with alignment following experience.

In addition, we analyzed the orientation discriminability of all L2/3 cell responses across latency (Fig. 4G). Orientation discriminability at the shortest latency was higher in experienced than naïve animals (p<0.001), consistent with increased reliability of feedforward inputs to L2/3 due to increased orientation discriminability in L4 units (Fig. 2B). While orientation discriminability increases slightly with latency in naïve animals (ANOVA effect of latency p<0.001), consistent with weak alignment driving weak amplification, this increase is larger in experienced animals (ANOVA latency-age interaction p<0.001), indicating improved alignment and amplification.

Together, our experimental findings regarding changes in coactivity between L4 neurons and L2/3 modular patterns and L2/3 intra-cellular orientation-tuning dynamics following the onset of visual experience can be explained, according to our model, by the maturation of two critical circuit properties: (1) the increase of orientation discriminability of L4 neurons providing inputs to L2/3, and a realignment of feedforward-recurrent interactions in L2/3 that enhances the selective amplification of orientation tuned signals.

## Discussion

These results show that early visual experience initiates a dynamic process of circuit reorganization that leads to enhanced feedforward-recurrent amplification supporting highly reliable sensory representations with interlaminar and temporal coherence. At eye opening, the naïve state of V1 circuits can achieve only poorly aligned feedforward-recurrent interactions in L2/3 in response to novel stimuli, such as oriented edges. These immature feedforward-recurrent interactions limit the reliability and temporal consistency of L2/3 responses at this early stage. Rapid changes in circuit organization following experience increase the reliability of feedforward L4-to-L2/3 inputs and transform the alignment of feedforward-recurrent interactions in L2/3 to amplify inputs elicited by the same orientation. These changes in input reliability and feedforward-recurrent alignment are both required to support the emergence of a highly reliable, temporally coherent modular representation of orientation.

Our findings provide new insights into the circuit mechanisms responsible for the initial state of the modular representation of orientation at the onset of experience^21–23^. A recent study has emphasized the high trial-to-trial variability in modular activity patterns that accompany the presentation of a single-oriented grating in visually naïve animals and the rapid increase in reliability and refinement of modular structure that follows experience^23^. Consistent with the predictions of a computational model, our results show that the modular responses to oriented stimuli seen in L2/3 at eye opening are the product of a combination of unreliable feedforward inputs from L4 and poorly aligned feedforward-recurrent interactions. But, despite poor alignment for orientation, we show that these naïve L2/3 recurrent circuits are amplifying (extracting) modular signals from highly variable input patterns supplied by L4 neurons. Thus, the initial modular map of orientation at the onset of visual experience is likely to reflect recurrent amplification of a subset of feedforward inputs that best resonate with the naïve state of recurrent connectivity.

While, in the mature cortex, evidence supports the view that feedforward inputs determine the basic tuning properties of neurons^40^, our intra-cellular data suggests a significant contribution of recurrent drive in the visual cortex at the onset of visual experience, improving the signal-to-noise ratio of novel incoming signals at this crucial stage in development. Continued recurrent amplification of visually driven input patterns combined with co-activity driven synaptic changes^41–43^ could enhance the selective amplification of tuning relative to variability in feedforward-recurrent interactions^8–12^. This process would lead to circuit changes, potentially involving refinement in afferent connections to L4, between L4 and L2/3, and within L2/3, that result in highly reliable representations^23^. Further experiments to probe the dynamic changes in feedforward-recurrent interactions chronically following eye opening are required to test these hypotheses further.

An organizational principle of cortical circuits based on experience-driven feedforward-recurrent alignment provides exceptional adaptability. The structure of stimulus-evoked feedforward input patterns reflects the properties of the animals’ environment. Thus, the development of aligned interactions between these inputs and recurrent connectivity results in an organized sensory representation whose foundations reflect the structure of the environment, as is observed experimentally^44–47^. This contrasts with the relative rigidity of some previous models, where modular sensory representations result from the experience-driven refinement of structural biases in feedforward circuits set up before eye opening^48–52^.

The primary visual cortex may not be unique in benefiting from an organizational principle based on feedforward-recurrent alignment. Recent studies have revealed modular spontaneous activity, possibly reflecting feedforward-recurrent interactions, in the early developmental stages of multiple areas, including diverse sensory modalities and those serving higher-order cognitive functions^34^. Therefore, the developmental sequence of alignment revealed here may represent a common organizational principle across cortical functions.

## Methods

### 1 In-vivo experiments

#### 1.1 Animal model and subject details

All experimental procedures were approved by the Max Planck Florida Institute for Neuroscience Institutional Animal Care and Use Committee and were performed in accordance with guidelines from the U.S. National Institute of Health. Only Female ferrets, procured from Marshal Farms, were used. Post-natal (P) age range was P27-31 for experiments in visually naïve animals and P36 or above and at least 5 days after eye opening for experiments in visually experienced animals. Juvenile ferrets were co-housed with jills on a 16-h light/8-h dark cycle.

#### 1.2 Surgical procedures

##### Intra-cranial virus injections

For experiments involving calcium imaging, viral injections were performed as previously described^1,2^. Briefly, we expressed GCaMP6s^3^ or JrGeco1a^4^ by microinjection of AAV2/1-hSyn-GCaMP6s/JrGeco1a-WPRE-SV40 (University of Pennsylvania Vector Core or Addgene®) into the visual cortex 1-4 weeks before imaging experiments. In developing ferrets, viral expression using the Synapsin promoter has previously been demonstrated to primarily label excitatory neurons^5^ and yield multiple millimeters of roughly uniform labeling^6^. Anesthesia induction was performed using either ketamine (50 mg/kg, IM) and/or isoflurane (1%–3%) delivered in O2 and then maintained with isofluorane (1%–2%). Atropine (0.2 mg/kg, IM) was administered to reduce secretions, while Buprenorphine (0.01 mg/kg, IM) and a 1:1 mixture of lidocaine and bupivacaine (injected directly into the scalp) were administered as analgesics. Animal temperatures were maintained around 36-38°C using a homeothermic heating blanket. When intubation was possible, animals were mechanically ventilated and end-tidal CO2 monitored. In all cases, heartrate was monitored throughout the surgery. Under aseptic surgical technique, a small craniotomy was made over visual cortex 4-7 mm lateral and 1-3 mm anterior to lambda. 1 ul of virus was pressure infused into the cortex through a pulled glass pipette at a single depth of 200-500 μm in up to 3 locations. This procedure reliably produced robust and widespread expression in excitatory neurons over an area that was at least 2 mm in diameter.

##### Recordings of visual responses

All animals were anesthetized and prepared for surgery as described above. Skin and muscle overlying visual cortex were reflected and a custom-designed metal headplate (8mm DIA) was implanted over the virus-injected region with MetaBond (Parkell Inc.). Then, a craniotomy and a subsequent durotomy were performed, and the underlying brain stabilized with agarose (2-3%). The headplate was covered with a thin 8 mm coverglass (Warner Instruments) with a drilled hole for access with electrophysiology probe or whole-cell glass pipette. Coverslip was fixed in place using a stainless-steel retaining ring (5/16’’ internal retaining ring, McMaster-Carr). Eye-lids were retracted to open the eyes and Phenylephrine (1.25%–5%) and tropicamide (0.5%) were applied to retract the nictitating membrane and dilate the pupil. A small amount of silicon oil was applied to the eyes to prevent eye drying and irritation. Prior to data collection, isoflurane levels were reduced from a surgical plane to .5-1.5%. After reaching a stable, anesthetic baseline (heart rate of 260-300 bpm), animals were paralyzed with pancuronium bromide (0.1-0.4 mg/kg/hr in lactated Ringer’s with 5% dextrose, delivered IV). After data acquisition was completed, isoflurane was raised to 5% and the animal was given Euthasol (0.5 ml, IV or IP) for euthanasia.

#### 1.3 Data acquisition

##### Calcium imaging

Animal was placed under epi-fluorescent microscope featuring Long-distance 4X Achrovid™ objective, Andor® Zyla 5.5 sCMOS camera, Thorlabs® LEDs, and EPI optical path setups. Images were acquired at 10-30Hz depending on calcium indicator expression levels but analyzed as 1 second temporal averages in all cases. Images covered an area approximately 6×6mm square and, after spatial down-sampling, yielded 515×512 pixels. Imaging data was analyzed only for pixels within the area of cortex that displayed strong expression of calcium indicator.

##### Extracellular electrophysiology

For extracellular electrophysiology, silicon probes were provided by Dr. Masmanidis at UCLA^7^. For simultaneous epi-fluorescence imaging and electrophysiology, linear probes with 32 or 64 channel spanning 920 or 1575 μm depth were used. For simultaneous epi-fluorescence imaging and electrophysiology, the probe was placed on V1 within the area of expression of calcium indicator using a micro-manipulator. For all experiments, the probe was driven down into the cortex until only the most superficial channels, or no channels, were out of the brain. Electrophysiology data was acquired using a Neuralynx® Digital Lynx 4SX system. For temporal alignment of electrophysiology data, imaging data, and stimulus, frame TTL triggers from the sCMOS camera and stimulus TTL triggers from the visual stimulation computer were recorded through the electrophysiology hardware.

##### In-vivo whole-cell

We followed similar methods for in-vivo whole-cell recordings as described in previous studies by our lab^2,8^. Briefly, recordings were performed by inserting a pipette through the hole of the coverglass into the agarose-filled craniotomy. A silver chloride reference electrode was inserted into the muscle. Recordings were made in the zero-current clamp mode. Pipette resistance was 5-8 MΩ. Internal solution contained (in mM) 110 K gluconate, 20 Na-Gluconate, 4 KCl, 10 HEPES, 10 K2-Phosphocreating, 4 Mg-ATP, 0.3 Na3GTP. pH was set to 7.2 and osmolality to ∼295 mOsm. Neurons were recorded from layer 2/3 (100 to 400 μm below the pia) using a Multiclamp 700B (Molecular Devices). Series resistance for recordings typically ranged from 20 to 80 MΩ. Analog signals were digitized using Spike2 (CED) and analyzed using custom-made MATLAB® based software.

#### 1.4 Visual stimulation

Visual stimuli were delivered on an LCD screen placed approximately 20–25 cm in front of the eyes using PsychoPy^9^. To evoke orientation responses, full-field square gratings at 100% contrast, 0.06 cycles per degree spatial frequency, and drifting at 3Hz temporal frequency were presented. 8 stimulus orientations and 16 corresponding drifting directions were sampled. In addition, ‘blank’ stimuli of 0% contrast were presented but not included in any analysis. Stimuli were randomly interleaved and presented for 1 s followed by 3-5 s of gray screen. 5-10 trials of each stimulus, including blank, were shown.

#### 1.5 Data analysis

##### Calcium imaging responses (Figure 1)

Epi-scale calcium responses were computed for each stimulation trial as:

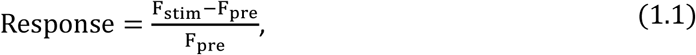

where F_stim_ is the temporally averaged fluorescence during visual stimuli (1 second long) and F_pre_ is the temporally averaged fluorescence during the previous second.

##### Extracellular electrophysiology (Figures 1-2)

Spike-detection, single-unit isolation, and response analysis were all done with custom MATLAB®-based software used in previous studies^10,11^. Spiking responses were computed for each stimulation trial as the difference in firing rate during stimulation (1 second) and that during the previous second. Single units for which no stimulus elicited a trial-averaged response above 2 Hz were deemed unresponsive and not included in any analysis.

##### Sink-source-analysis of local-field potentials for laminar idenificaion (Figure S1)

To identify laminar borders in our multi-channel electrophysiology data we first computed local-field-potentials (LFP) in response to either full-screen changes in luminance or m-sequence noise stimuli. LFP responses were then band-pass filtered (5-50 Hz), which reliably yielded clean signals in our recordings. To reduce noise, channel whitening was performed on the filtered LFP data. Current source density was then calculated by the inverse method^12^ using Matlab® code made public by Timothy Olsen (Timothy Olsen (2024). Current source density (CSD), MATLAB Central File Exchange). We then identified the laminar borders between layer 2/3, layer 4, and layers 5/6 based on sink-source transitions, as described in previous studies^13–15^.

##### Unit-to-modular coactivity (Figure 1)

Only imaging pixels corresponding to an area within a 1mm distance from the electrode insertion site were considered for this analysis. This allowed us to assess correlations between single-unit spiking and modular calcium responses across a similar distance for different experiments where the spread of calcium indicator expression varies. In addition, pixels within 250 μm of the insertion location were discarded to disregard local-circuit activity correlations and focus on correlations with neighboring functional modules.

Due to both imaging and spiking data not displaying normal distributions, we decided to compute a non-parametric estimator of coactivity. First, calcium responses were band-passed Gaussian-filtered across space (20-200 μm/cycle) and Z-scored across pixels for each trial. This step allowed us to remove any correlations due to the overall responsiveness of the cortex, revealing correlations with modular activity patterns. Then, for each cell recorded, we computed a trial-to-trial spike-weighted average of the calcium response. Next, we computed a shufie distribution of this spike-weighted average by randomly re-assigning spiking responses across trials. 100 iterations of this randomization were made to compute the distribution. Finally, the non-shufied spike-weighted average was z-scored by the shufie distribution yielding the final unit-to-modular coactivity. Thus, the sign of this coactivity quantification represents the sign of the correlation while the absolute amplitude represents the deviation from a randomized expectation for each experiment and single-unit. The strength of unit-to-modular coactivity was computed as the mean absolute value of the non-parametric correlations across imaging pixels. Due to the Z-score normalization of the correlation, the coactivity strength expected by chance is the mean of a half-normal distribution (J ∼ 0.8).

##### Unit-to-modular orientation correspondence (Figure 1)

First, we aimed to calculate the correlation between the orientation tuning of each unit’s spiking responses and that of the modular calcium response of each pixel in the imaging data, i.e., co-tuning. To prevent noise correlations from contaminating our estimation of co-tuning, we computed, for each unit and pixel, two tuning curves by averaging responses across even or odd-numbered trials. We then correlated unit and pixel tuning curves computed using complementary trials. Note that this approach yields two estimations of tuning correlation for each unit-pixel pair, which we averaged to compute the final quantification of unit-to-pixel co-tuning.

Next, for each unit, we correlated its co-tuning and coactivity with pixels across the cortical surface. High correlations indicate that L2/3 modules with high trial-to-trial coactivity with a unit display co-tuning with the unit, i.e., orientation correspondence.

##### Orientation preference mismatch between units and coactive modules (Figure S2)

To test if our results regarding unit-to-modular coactivity orientation correspondence were confounded by uncertainty in our estimation of units’ orientation preference due to trial-to-trial response variability, we first quantified such uncertainty as the mean absolute error in the estimation of units’ preference across 100 randomized re-sampling (with replacement) of single-trial responses.

The expected impact of uncertainty on the quantification of orientation correspondence described above is not clear or easily quantified. Considering this, we instead calculated the orientation preference mismatch between each unit spiking responses and their coactive modular pattern. This simpler metric of orientation correspondence can be directly compared with orientation uncertainty, and they should be similar if the latter fully explains the former.

The orientation preference of units’ spiking and pixels’ calcium responses was computed as their circular mean across stimuli following the formula:

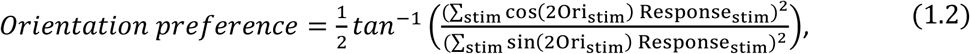

where Ori is the orientation of the stimulus.

The orientation preference of each unit’s coactive modular pattern was then calculated as the coactivity-weighted circular mean across pixels using the formula:

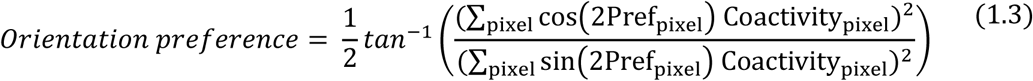

We then calculated the circular distance between units and their coactive modular pattern preferred orientations and compared it to the uncertainty of units’ orientation preference estimation.

##### Single-unit orientation discriminability (Figure 2)

This was quantified as Cohen’s D between trial-to-trial responses to the preferred grating stimulus, defined as that eliciting maximum trial-averaged response (considering both grating orientation and direction), and the two orthogonal gratings (moving in opposite directions). The equation was:

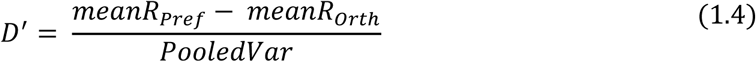

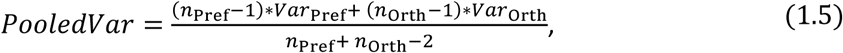

were *meanR* is the mean response across trials, *Var* the trial-to-trial variance, *Pref* represents the preferred grating orientation and direction, and *Orth* represent the orthogonal gratings moving in either direction.

##### Whole-cell sub-threshold responses (Figure 4)

First, we computed the response latency of each neuron as the time when its membrane potential increased significantly from that during the 10 ms prior to stimulus onset, considering data across all trials and stimuli. Statistical significance was defined as p < 0.001 using paired t-tests. After correcting sub-threshold responses by this latency, we computed a full response for each trial as the difference in membrane potential between a one-second period right after response onset and a one-second period preceding response onset. Next, we computed responses for different latencies as the difference in membrane potential between the 1 s preceding response onset and a 50 ms temporal window centered at variable intervals from stimulus onset, ranging from 0 to 1 s in steps of 10 ms. For responses at all latencies and the full response, orientation preference and discriminability were computed using the same formulas described previously for single-unit spiking responses (Eq. 1.2 and 1.4).

#### 1.6 Statistics

For all comparisons of data involving spiking responses (which do not follow a normal distribution), we used non-parametric Wilcoxon rank-sum tests. This included all comparisons in Figures 1-2.

For all comparisons of data involving sub-threshold whole-cell responses shown in Figure 4, we used parametric tests including t-tests and analysis of variance (ANOVA).

For all data, all tests were two-tailed and the replicate was the single-unit (figures 1-2) or cell (figure 4). The number (N) of animals in our experiments was; 6 naïve and 5 experienced for data in Figures 1-2; and 20 naïve and 12 experienced for data in Figure 4.

#### 1.7 Data availability statement

The data and code that support the findings of this study are available from the corresponding author upon reasonable request.

### 2 Computational modeling

#### 2.1 Motivation

The goal of our model was to examine how the reliability of feedforward inputs from layer 4 (*L*4) to layer 2/3 (*L*2/3) together with the alignment of these inputs with the recurrent network in *L*2/3 determine several important cortical network response properties characterizing orientation selective responses in the visually experienced cortex (Figures 3, 4A, and 4B). These response properties are interlaminar coactivity strength and orientation correspondence (Figure 1), orientation discriminability (Figure 2) and the consistency of orientation tuning during sustained stimulation (Figures 4C-G).

To this end, we modeled *L*2/3 employing a recurrent network of rate units, whose properties were constrained by replicating key properties of *L*2/3 spontaneous activity prior to eye opening in ferret visual cortex^6,16^. To model visual grating-evoked activity in *L*2/3 after eye opening, we analyzed the responses of this network to orientation-selective feedforward inputs from *L*4, varying systematically their degree of reliability (orientation discriminability) and alignment with the recurrent network. Our approach is consistent with the observation that both visual-grating evoked and spontaneous activity exhibit a modular organization which shares basic structural attributes including a similar spatial wavelength estimated as 0.75 - 1.0 *mm*^16,17^.

In the following, we define the dynamics of the network model (2.2), describe the structure of its recurrent connectivity (2.3) and of its orientation-tuned feedforward inputs (2.4), and finally describe its implementation and data analyses that yielded the results presented in the main text (2.5).

#### 2.2 Network model

Building on our previous work^6,18^ we modeled the neuronal modular activity of *L*2/3 using a standard recurrent network of a single population of rate units^19,20^

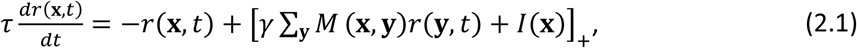

where *r*(**x**, *t*) is the activity of a unit at position **x** at time *t*, τ is the neuronal timescale, [*x*]_+_ = max{0, *x*} is the neuronal activation function, γ is a factor controlling the overall strength of recurrent connections, *M*(**x**, **y**) is the weight of the recurrent connection from the unit at location **y** to the unit at location **x**, and *I*(**x**) is the external feedforward input to the unit at location **x**, which is assumed constant in time for simplicity.

#### 2.3 Recurrent connectivity

The recurrent connectivity of our model was constructed to reproduce two core features of early spontaneous activity in *L*2/3 ferret visual cortex prior to eye opening: its modular structure and its moderately low-dimensional organization on the scale of several millimeters, giving rise to long-range correlations^6^.

To reproduce the modular structure of cortical activity, we chose a recurrent connectivity with the spatial profile of local excitation / lateral inhibition (LE/LI), which is known to provide a simple, yet robust dynamical mechanism for modular cortical activity, and for which a recent study provides empirical evidence^18^. The concrete implementation of the LE/LI mechanism we used is a difference-of-Gaussians connectivity

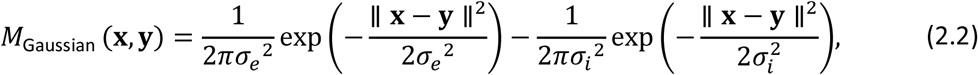

where σ_e_and σ_i_ control the range of local excitation and lateral inhibition, respectively. For such isotropic LE/LI-connectivity the eigenvectors of *M* are plane waves, and the spectrum is peaked at a wavenumber *k*_0_ = 2π/Λ with the typical spatial scale Λ given by

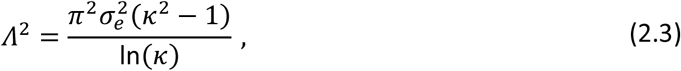

where *k* = σ_i_/σ_e_ is the ratio of the spatial range of inhibitory and excitatory connectivity (set equal to 2).

To account for the moderately low-dimensional organization of spontaneous activity, we assumed a moderate degree of heterogeneity of LE/LI interactions across cortex. Such heterogeneity captures possibly variations of connectivity across cortical domains and was found in previous work to have a strong impact on the dimensionality of activity patterns^6,21^. At eye opening, the structure of this heterogeneity may reflect the properties of the correlation structure of early spontaneous activity. Therefore, we constructed the recurrent connectivity similar to^18^ by adding a heterogeneous perturbation *M*_het_ (**x**, **y**) to the LE/LI connectivity,

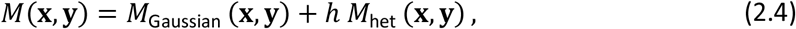

where ℎ controls the strength of heterogeneity. This heterogeneous part of the connectivity is defined by

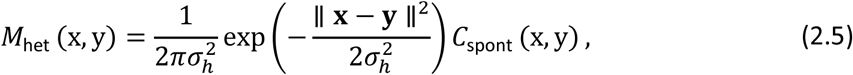

where σ_m_ controls the range of modular connectivity (set to the same value as σ_i_), while *C*_spont_ (x, y) mimics the spontaneous activity correlation structure of early spontaneous activity. This correlation structure was modeled as

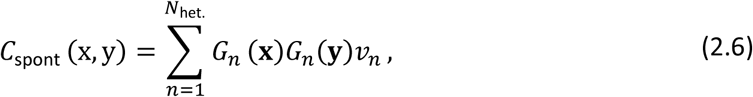

using random modular patterns *G_n_*, constructed by applying a band-pass filter centered around *k*_O_ to Gaussian white noise patterns and then orthonormalizing them. The *v_n_* are exponentially decaying weights defined as

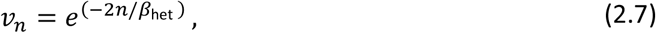

where β_het_ determines the dimensionality associated with the correlation *C*_spont_ ^22^, which we set equal to 10 to match roughly the dimensionality of spontaneous modular activity observed experimentally^6,16,22^.

We rescaled the inputs to each network unit to have zero mean and uniform variance, consistent with homeostatic mechanisms balancing the input between excitation and inhibition, equally across cortex. To achieve this, we first normalized for each location **x** all negative recurrent weights from **y** to **x**, such that the sum of recurrent weights to location **x** equals zero. Then, we normalized for each location **x** all recurrent weights from **y** to **x**, such that the standard deviation of recurrent weights to location **x** equals 1. For further analysis, *M* was scaled such that the real part of its maximal eigenvalue is equal to 1. Finally, the two model parameters ℎ and γ were chosen to reproduce the experimentally observed dimensionality^6,16,22^ when driving the network with spatially unstructured random inputs. To this end, we simulated 100 responses to unstructured input patterns constructed according to

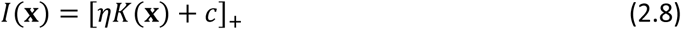

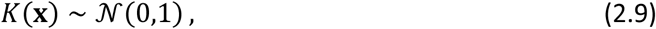

where *c* is a constant activation, set to 0.1, and η controls the standard deviation (modulation) of the distribution of inputs, set to 0.01. Input patterns *K*(**x**) were low-passed filtered (Gaussian filter kernel σ_Low_ = 3) to model overlapping pools of feedforward connections across neighboring *L*2/3 units. In addition, a rectification was applied to suppress negative inputs (affecting < 0.001% of all inputs). Finally, input to each location was normalized across trials to zero mean and unit standard deviation to avoid biases in the overall feedforward input drive across cells. To achieve a better comparison with experiment, model responses were band-pass filtered using a difference of Gaussian filter defined by σ_e_ and σ_i_ before calculating the dimensionality as participation ratio^23^. Based on the results from these simulations and analysis (Fig. S4), the parameters ℎ and γ were set to 0.5 and 1.05, respectively.

#### 2.4 Orientation-tuned feedforward inputs

We assumed orientation-selective feedforward input to each network unit, varying in reliability across units. The input to a given unit was assumed to represent the aggregate feedforward inputs received by a *L*2/3 neuron from a local pool of *L*4 neurons. The total input drive to a unit at location **x**, when presented with stimulus θ on trial *i*, was modeled as

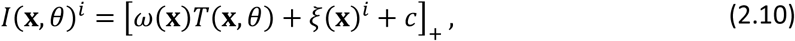

where *T*(**x**, θ) is the tuned component of the input, ω(**x**) controls its reliability, ξ(**x**, tr) describes a trial-dependent noise component, and *c* is the same constant activation function as in equation (2.8). In addition, a rectification was applied to suppress negative inputs (affecting < 0.001% of all inputs). The tuned input component for grating orientation θ was described by

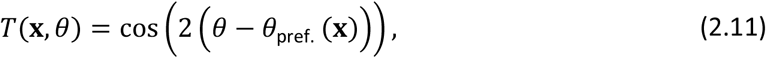

where θ is the stimulus orientation and θ_pref._ (**x**) the preferred orientation of the input to the unit at location **x**. In the following, we describe how the input reliability ω(**x**) and the input noise ξ(**x**)^i^ are constructed.

##### Input reliability

Rather than providing all units with inputs displaying equal levels of reliability, we considered variable input reliabilities across units, as this is expected to reflect more closely the diversity of response reliability across *L*4 neurons at any given developmental stage (see Figures 2B and 3C). To this end, we sampled reliabilities according to

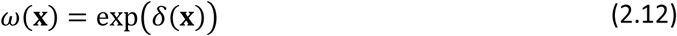

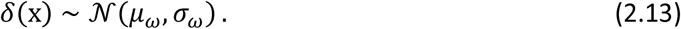

The parameters μ_w_ and σ_w_ control the median and spread of input reliability across units and were fit to the distribution of orientation discriminability of *L*4 single-unit responses in our experimental data (Fig. 3C).

##### Input noise

We found that for our model to produce input-to-network coactivity with poor orientation correspondence, like that observed for interlaminar coactivity in naïve animals (Fig. 1F), the input noise in the model must be correlated between units across trials (Fig. S7). To produce such correlations, we modeled the noise component of the input as

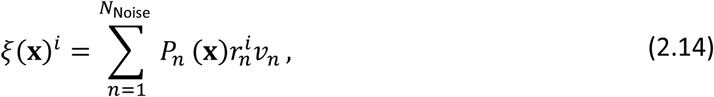

where *P_n_* are Gaussian white noise patterns with zero mean and unit standard deviation, *r*^i^ are scalars sampled independently from a normal distribution for each field *n* and trial *i*, and

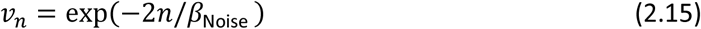

are weights applied to each random field to control the dimensionality of ξ(**x**)^i^ and thus the degree of correlations across inputs.

We found that moderately large values of β_Noise_ (we used β_Noise_ = 67) produce interlaminar coactivity for small alignment and low input reliability similar to that measured experimentally between *L*4 units and modular *L*2/3 response patterns in naïve animals (Fig. 1F). As above for the unstructured inputs used for modeling spontaneous activity, the input noise patterns ξ(**x**)^i^ were low-passed filter across space (Gaussian filter kernel σ_Low_ = 3). Input noise to each cell was then normalized across trials to zero mean and unit standard deviation.

Note that this construction of inputs implies that the discriminability of inputs, *D*′(**x**), is linearly related to ω(**x**) and expressed through

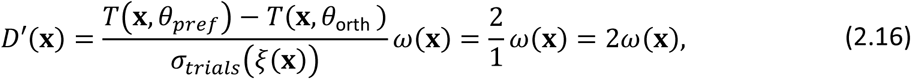

where σ_trials_(ξ(**x**)) is the standard deviation of noise across trials and *T*(**x**, θ_Pref_) and *T*(**x**, θ_Orth_) are the tuning components at the preferred and orthogonal orientations, respectively.

Finally, we normalized ω(**x**)*T*(**x**, θ) + ξ(**x**)^i^, by its standard deviation across all stimuli and trials, which has the effect that variance across all trials and stimuli is the same for each unit. We then rescaled this location dependent part in equation 2.10 by a factor η = 0.01, such that its standard deviation is 10% of the spatially uniform part of the input *c*.

These parameter settings resulted in a modulation of total (feedforward and recurrent) inputs received by modeled units in the range from 0.3-0.75 depending on input reliability and alignment (mean of 0.53). This was similar to the range of modulation of inputs received by *L*2/3 neurons in our in-vivo whole-cell data (mean Naïve: 0.42, Experienced: 0.64).

##### Feedforward-recurrent alignment

To systematically study the impact of feedforward-recurrent alignment on network response properties, we varied the spatial layout of the tuned input component *T*(**x**). To this end, we interpolated the orientation preference of input to location **x** as

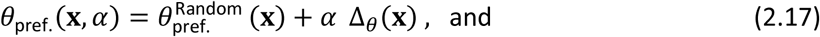

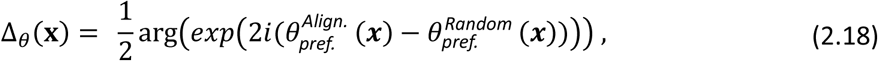

where θ^Random^_pref._ is a random layout, in which orientation preferences are sampled from a uniform distribution [0, π], to which a fraction α of the difference Δ_0_(**x**) between the aligned θ*^Align.^_pref._* (***x***) and random layout is added. Here, α acts as interpolation parameter, so that for α = 0 and α = 1 the orientation preference θ_pref._(**x**, α) is identical to the random and aligned layout, respectively. As one possibility to generate a relatively well-aligned state, we used

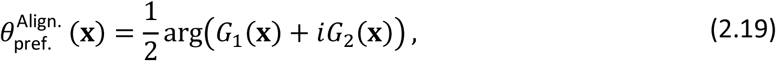

where *G*_1_(**x**) and *G*_2_(**x**) are the two modular random field patterns with strongest weights used to produce the heterogeneity in the recurrent connectivity (see Eq. 2.6). We defined the effective feedforward-recurrent alignment as

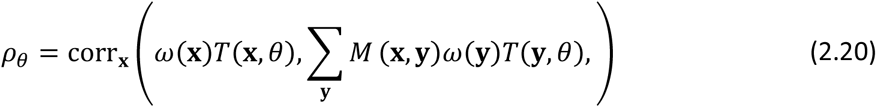

based on the tuned component of the input ω(**x**)*T*(**x**, θ) and the recurrent connectivity *M*. Note that this defintion of feedforward-recurrent alignment extends the definition used in our previous study^22^ making it applicable also to non-symmetric matrices.

#### 2.5 Implementation and analysis

We implemented the network with 60 X 60 units placed on a regularly-spaced grid, using periodic boundary conditions. This grid was considered to represent 4 X 4 mm of cortical surface (grid inter-unit distance Δx corresponding to 66.7 μm).

For the LE/LI connectivity, we used σ_e_ = 1.8 and σ_i_ = 3.6, implying a wavelength of approximately Λ ∼ 12 (in Δx) or 0.8 *mm*.

We simulated activity in the network for a period of *T*_total_ = 60τ, which was selected to allow the network to reach an activity pattern near steady state, defined as > 0.995 correlation between activity patterns separated by 10τ, under all input conditions. We numerically integrated the dynamics using a fourth-order Runge-Kutta method from *t* = 0 to *t* = *T*_total_ using a time step *dt* = 0.15τ. For all simulations we used the initial conditions *r*(**x**, *t* = 0) = 0.

Similar to the empirical data, for each pair of input reliability and alignment, we simulated responses for 8 equally spaced orientations, and 10 trials for each orientation, amounting to 80 simulations.

All results were averaged over 10 different samples of recurrent networks using different samples of modular field patterns (*G*_n_) underlying recurrent connectivity. For each network, we simulated responses elicited by inputs with varying degrees of feedforward-recurrent alignment and reliability distributions. Results in Figures 3E-G, 4B, and S7A were averaged across units/inputs and across the 10 recurrent networks. For each recurrent network, we randomly sub-sampled 20% of units/inputs when computing response properties. Data for individual recurrent networks are shown in Figures S6 and S7B-C for a subset of input reliability and alignment conditions.

##### Varying input reliability

To systematically study how the network properties depend on the reliability of inputs (Figure 3E-G), we estimated the parameters μ_w_ and σ_w_ (see eq. 2.13) in the visually naive and experienced cortex by matching the distribution of orientation discriminability defined in Eq. 2.16 to the distribution of orientation discriminability of *L*4 single-unit responses, and then linearly interpolated these parameter values to arrive at estimates of input reliability distribution in between these two developmental time points (Fig. 3C). Thus, low and high input reliability conditions (1 and 7 in Figures 3E-G, S7A, and 4B) were constructed by fitting μ_w_ and σ_w_ to match the distributions of input orientation discriminability with that measured experimentally in L4 single-units in naïve or experienced animals, respectively (Fig. 3C). Intermediate reliability levels were constructed by linear interpolation of μ_w_ and σ_w_. Orientation discriminability distributions in *L*4 single-units were not statistically different from that in short-latency (< 100 ms) inputs recorded through whole-cell recording at each age (rank-sum p values; Naïve: 0.73. Experienced: 0.22), indicating it is a reasonable estimate of the discriminability of aggregated feedforward inputs to *L*2/3.

##### Varying feedforward-recurrent alignment

We varied α to study different degrees of feedforward-recurrent alignment and for each α averaged ρ_0_ over all simulated angles θ and randomized recurrent connectivity instances. Note that no input reached an effective alignment of 1, as this would require spatially restricted input matching local recurrent connectivity. For a given level of α, effective alignment varied slightly across the 10 randomized recurrent networks. However, it was always within ±0.05 of the target value. In heat map plots (Fig. 3E-G, 4B, and S7A) columns represent data for 10 α values selected to target effective alignments from 0 to 0.9 in steps of 0.1. Mean effective alignment across randomized recurrent networks for each level of α is reported on Table 4.

##### Analysis of coactivity

One goal of our model was to assess the effects of input reliability and alignment on the interlaminar coactivity between *L*4 units and modular responses in the *L*2/3 recurrent network. For simplicity, we did not model individual *L*4 neurons and their connectivity structure to *L*2/3. Instead, we used coactivity between feed-forward input received by individual *L*2/3 units and modular activity in the modeled *L*2/3 network to probe the effects of input reliability and alignment on interlaminar coactivity. The strength and orientation correspondence of input-to-network (Fig. 3F-G) and unit-to-network (Fig. S7) coactivity were computed as described for unit-to-modular coactivity of experimental data (Fig. 1C and 1F). Coactivity was only considered for distances matching those used for experimental data (0.25-1 mm).

#### 2.6 Tables of model parameters

See Tables 1-4 for a summary of parameters. All networks were simulated using custom-written code in Python. Analysis of modeling results was done using custom-written code in Matlab®. Code underlying the model will be made available in GitHub upon publication.

**Table 1:** Network spatial and dynamics parameters.

| Symbol | Parameter | Value |
| --- | --- | --- |
| | Grid size | $60 \times 60$ |
| | | $4 \times 4 \text{ mm}$ |
| $\Delta x$ | Inter-unit distance | 1 |
| | Conversion factor for inter-unit distance | $66.7 \mu m$ |
| $T_{\text{total}}$ | Total simulation time | $60 \tau$ |
| $dt$ | Timestep | $0.15 \tau$ |
| $\tau$ | Neuronal time constant | 1 |

**Table 2:** Recurrent connectivity parameters.

| Symbol | Parameter | Value |
| --- | --- | --- |
| $\sigma_e, \sigma_i$ | Range of LE/LI connectivity | 1.8, 3.6 |
| $\sigma_h$ | Range of heterogeneous connectivity | 3.6 |
| $\beta_{\text{het.}}$ | Dimensionality of heterogenous connectivity | 10 |
| $N_{\text{het.}}$ | Number of gaussian fields ($G_n$) for constructing heterogeneous connectivity | 30 |
| $h$ | Strength of heterogeneity | 0.5 |
| $\gamma$ | Recurrent connectivity strength | 1.05 |

**Table 3:** Input parameters.

| Symbol | Parameter | Value |
| --- | --- | --- |
| $c$ | Constant activation | 0.1 |
| $\eta$ | Input standard deviation across all trials and stimuli | 0.01 |
| $\beta_{\text{Noise}}$ | Noise dimensionality for structured inputs | 67 |
| $N_{\text{Noise}}$ | Number of noise patterns for structured inputs | 100 |
| $\sigma_{\text{Low}}$ | Gaussian filter kernel for low-pass filter | 3 |

**Table 4:**
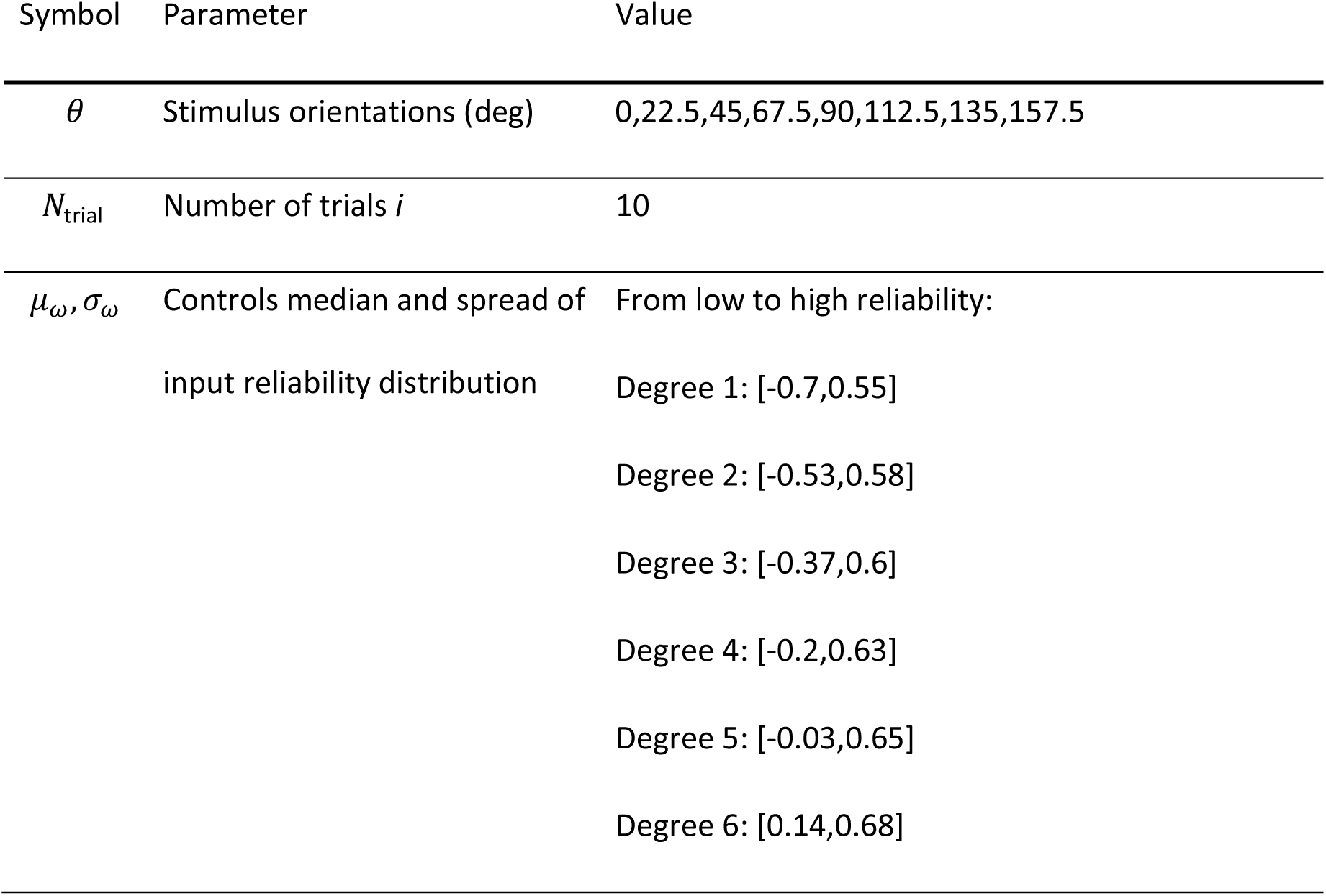
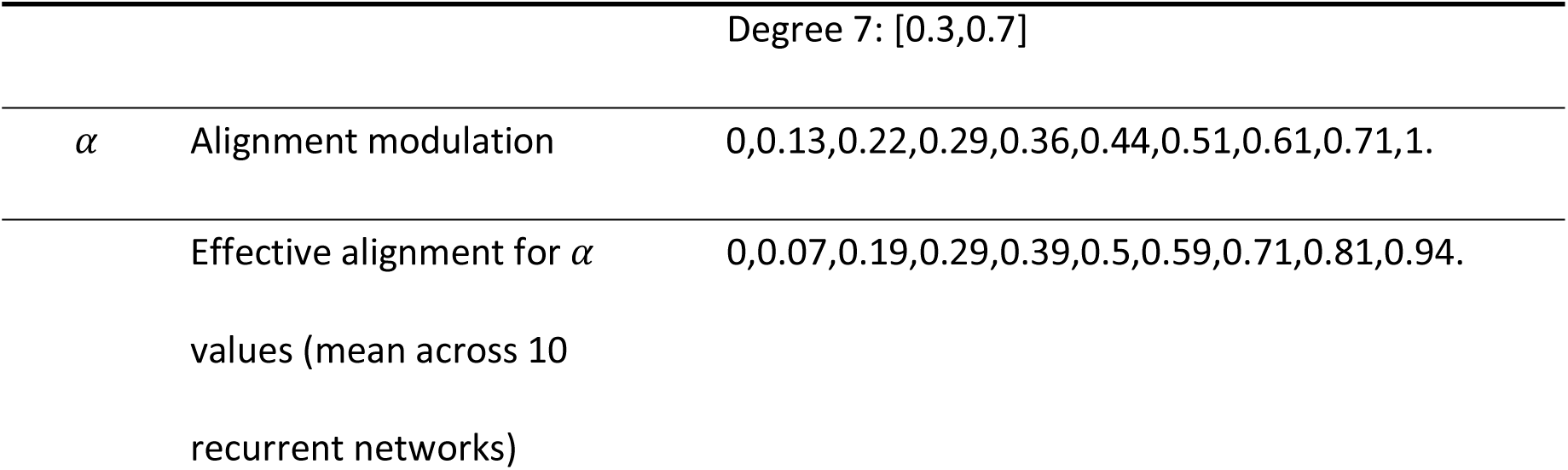
Simulation parameters.

## Acknowledgements

National eye institute funding: K99EY034936-01A (A.A.L.), 2R01EY011488 (D.F.); LOEWE Focus Center for Multiscale Modelling in Life Sciences (CMMS) (M.K, S.T.), the International Max Planck Research School for Neural Circuits in Frankfurt (S.T.), SPP 2041 “Computational Connectomics” (S.T.). The funders had no role in study design, data collection and analysis, decision to publish, or preparation of the manuscript.

We would like to thank G. Kreal and J. Drayer for technical assistance, as well as members of the Fitzpatrick and Kaschube laboratories for helpful discussions.

## Author contributions

Experimental procedures for electrophysiology and calcium imaging and associated data analysis were performed by Augusto Abel Lempel and Clara Tepohl under the guidance and mentorship of David Fitzpatrick. Design, implementation and analysis of recurrent network model and associated data analysis were performed by Augusto Abel Lempel and Sigrid Trägenap under the guidance and mentorship of Matthias Kaschube. Augusto Abel Lempel, Sigrid Trägenap, Matthias Kaschube, and David Fitzpatrick all contributed to manuscript preparation and data presentation design.

## Additional information

All authors declare no competing interests.

## Supplemental figures

**Figure S1:**
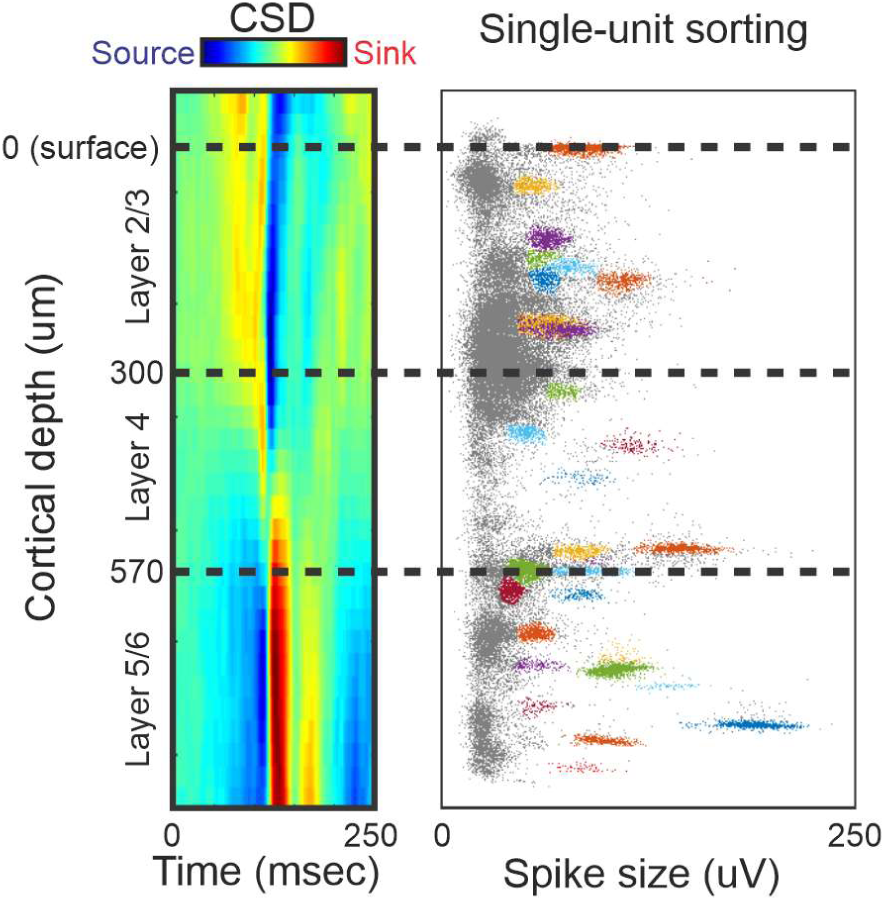
Identification of laminae in multi-channel electrophysiology experiments using sink-source analysis of local field potentials. The heat map on the left shows current source density across probe channels at different cortical depths and across time following a full-field luminance change stimulus. The scatter plot on the right shows the depth and size of spikes detected along the probe. Colors indicate single-unit activity sorted using custom-made software. Dashed lines indicate laminar transitions identified by sink-source transitions in the current source density profile.

**Figure S2:**
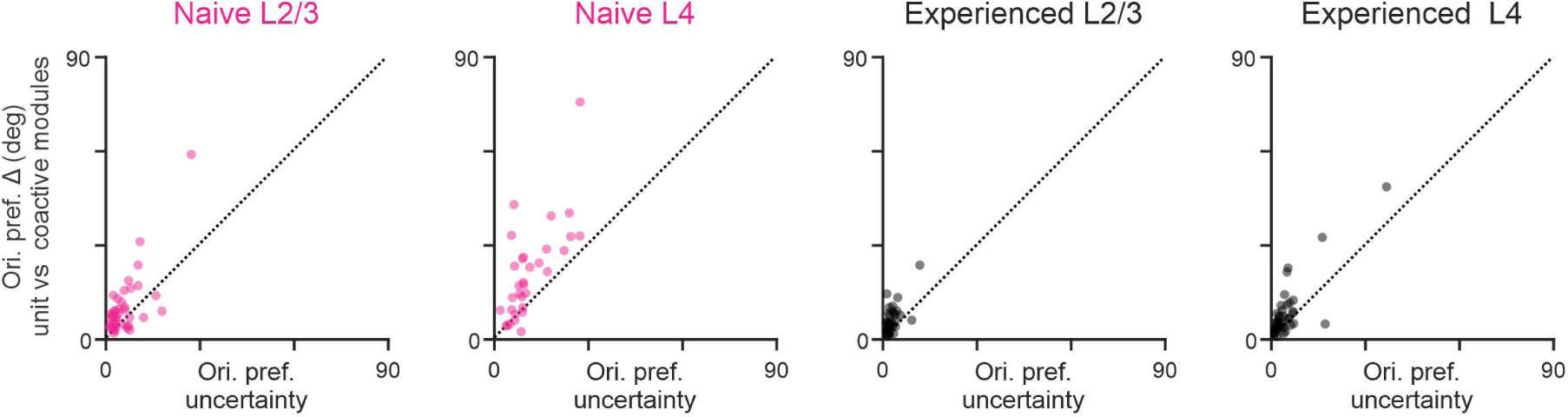
Orientation preference mismatch between units and coactive modules is not explained by poor accuracy of orientation preference estimation due to trial-to-trial variability in the units. Scatter plots show, for each unit, its orientation preference mismatch (Δ) to coactive layer 2/3 (L2/3) modules and the uncertainty in the estimation of its orientation preference. The dotted line represents a mismatch equal to estimation uncertainty. Many naïve layer 4 (L4) units display a preference mismatch to coactive modules that is considerably higher than the uncertainty in their orientation preference. Uncertainty was computed as the mean error in estimated orientation preference across 100 random re-samplings (with replacement) of single-trial data.

**Figure S3:**
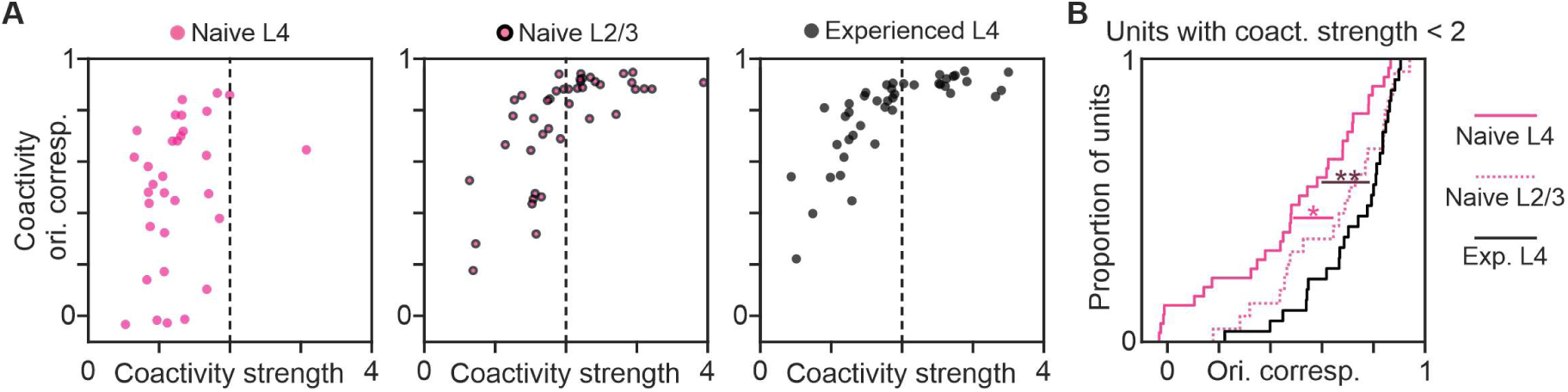
Independence of coactivity strength and orientation correspondence for naïve layer 4 (L4) units. **A** - Scatter plots of orientation correspondence versus unit-to-modular coactivity strength for naïve L4 units, naïve layer 2/3 (L2/3) units, and experienced L4 units. **B** - Cumulative density function plot of coactivity orientation correspondence for units with weak coactivity (strength below 2). Naïve L2/3 and experienced L4 units with weak coactivity display higher orientation correspondence than naïve L4 units with weak coactivity. *: p < 0.05, **: p < 0.01 using Wilcoxon rank-sum test.

**Figure S4:**
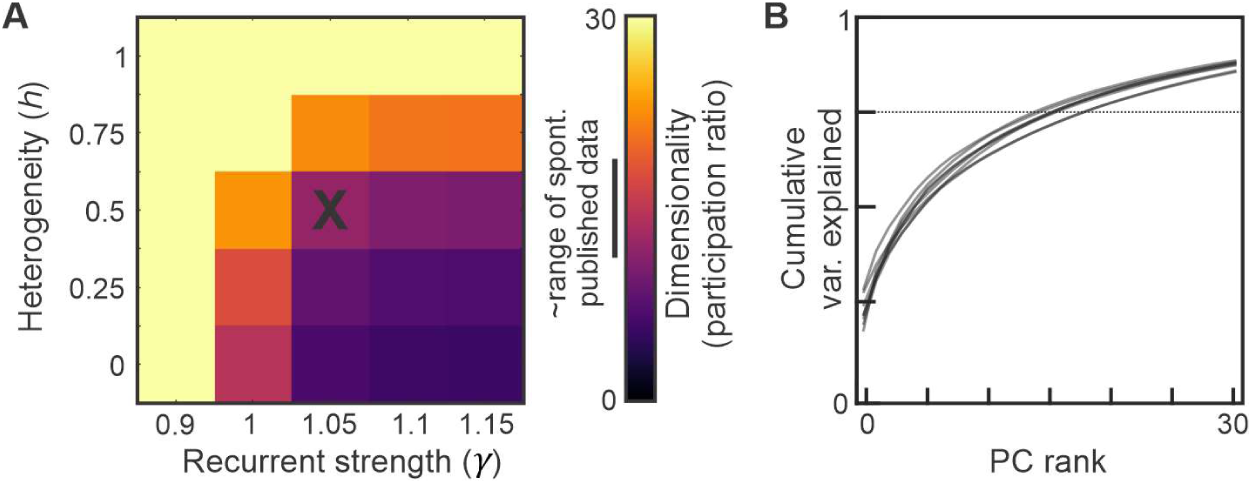
Recurrent connectivity model parameters were chosen to reproduce the dimensionality of spontaneous activity of layer 2/3 in published data. **A** - Heat-map of the dimensionality of responses elicited by spatially weakly smoothed unstructured input in the modeled recurrent network, quantified as the participation ratio (see Methods), as a function of recurrent connectivity strength (γ) and heterogeneity (h). Bar to the left of the scale-bar represents the range of published data^6,16,22^. Black X represents selected model parameter values used for results shown in main figures (γ: 1.05, h: 0.5). Data in this graph were averaged across 10 randomly generated instances of recurrent networks. B - Cumulative variance explained as a function of principal component (PC) rank. This data is for model parameter values used for results shown in main Figures 3 and 4 (γ: 1.05, h: 1); each line corresponds to a randomly generated instance of recurrent connectivity.

**Figure S5:**
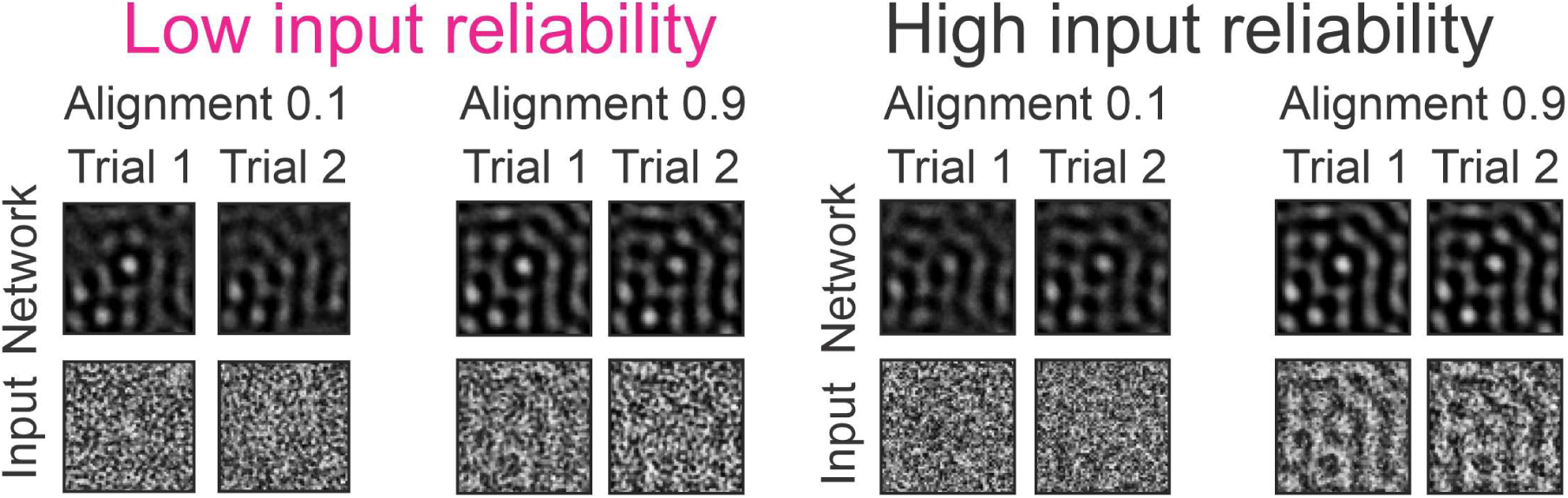
The network model produces modular responses when elicited by orientation-tuned, feedforward inputs. Examples of feedforward input and network activity patterns for low and high levels of input reliability and feedforward-recurrent alignment. For each level of reliability and alignment, two example trials are shown to indicate the degree of variability in the response patterns, which is strongest for low reliability and alignment. White indicates maximal, black zero activity (input) in each panel. Network responses are shown at the end of the simulated stimulation period (60T).

**Figure S6:**
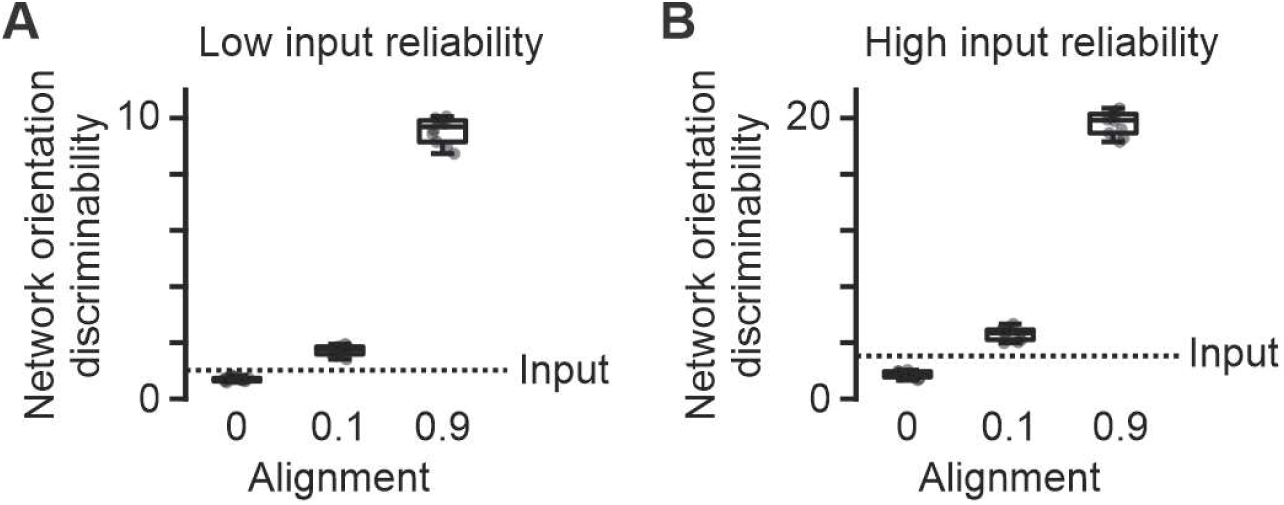
Feedforward-recurrent alignment is required for the amplification of orientation discriminability. Plots show the orientation discriminability of recurrent network responses elicited by inputs with low (**A**) and high (**B**) reliability under different levels of feedforward-recurrent alignment. Each dot represents average discriminability across units for a single instance of recurrent connectivity. Box plots show data distribution across recurrent connectivity instances (N=10). The dotted lines represent the mean orientation discriminability of external feedforward inputs. Recurrent network responses display higher discriminability than inputs only when alignment is greater than 0.

**Figure S7:**
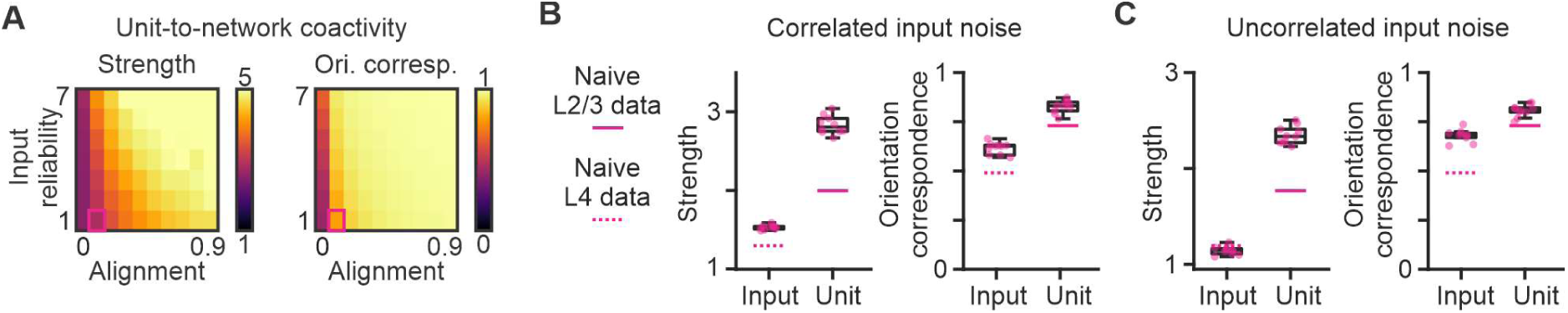
In the model, unit-to-network coactivity is stronger than input-to-network coactivity, and displays also a higher orientation correspondence, when inputs have poor reliability, poor alignment, and correlated noise. **A -** Heat maps of unit-to-network strength and orientation correspondence as a function of input alignment and reliability in the model. The pink box indicates input properties for data displayed in B. **B -** Input-to-network (input) and unit-to-network (unit) coactivity strength and orientation correspondence under low (0.1) input alignment and reliability (compare Fig. 4F and 4G). Each pink dot represents average discriminability across units for a single instance of recurrent connectivity. Box plots show data distribution across recurrent connectivity instances (N=10). For these simulations, external feedforward inputs displayed correlated trial-to-trial variability (noise) across model units. Pink lines represent mean unit-to-modular coactivity strength and orientation correspondence for layer 2/3 (solid) and layer 4 units (dotted) recorded in naïve animals (reproduced from Figure 1C,F). Note that, while orientation correspondence is slightly higher for model than experimental data, the difference in its mean between input and unit (0.21) is similar to that between layer 4 and layer 2/3 units (0.27). **C –** Same as B but for external inputs displaying no noise correlation across modeled units. Note that input-to-network orientation correspondence is higher compared to the model with correlated input noise, and its difference to the unit-to-network value is considerably reduced (0.12).

**Figure S8:**
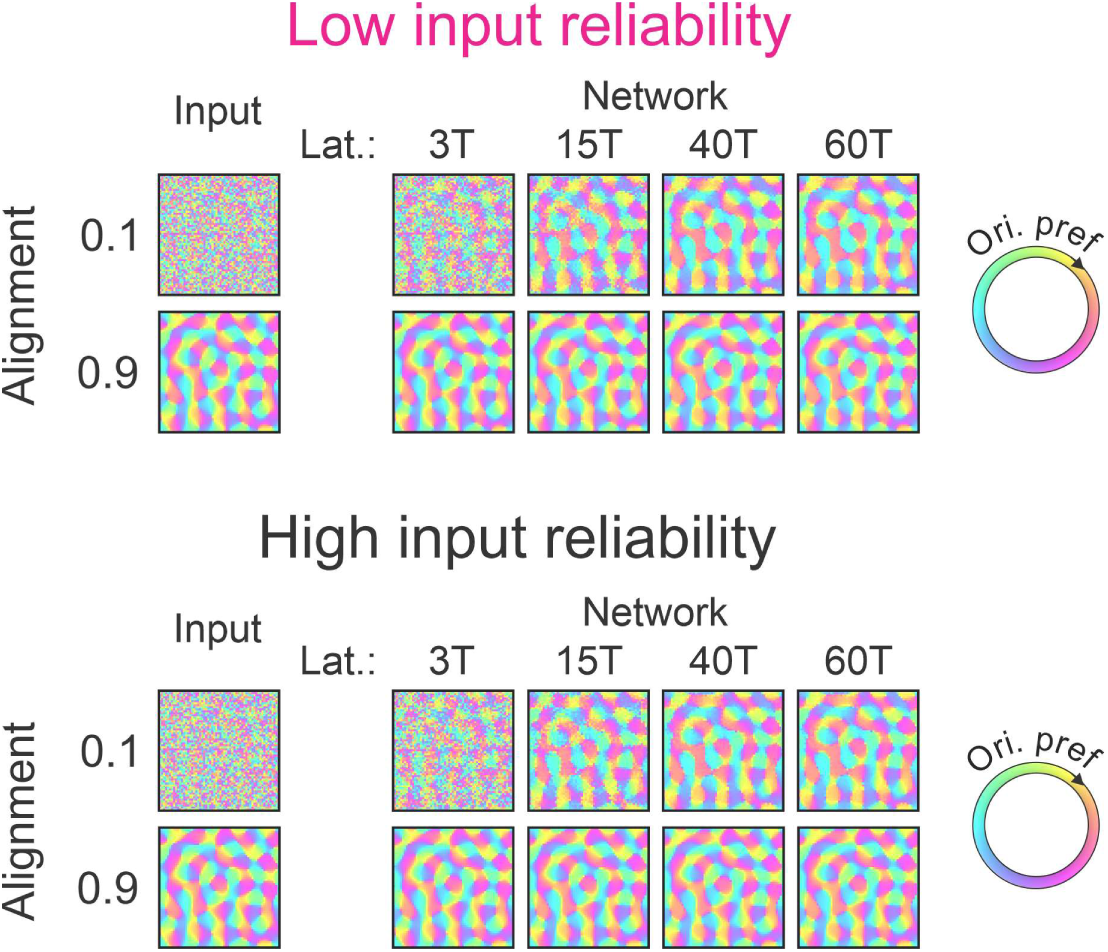
Because of its recurrent network, the model displays a well-organized modular orientation map for sustained stimulation, even when receiving inputs with low reliability (top). However, for weak feedforward-recurrent alignment (0.1), many units shift their orientation preference after stimulus onset.

**Figure S9:**
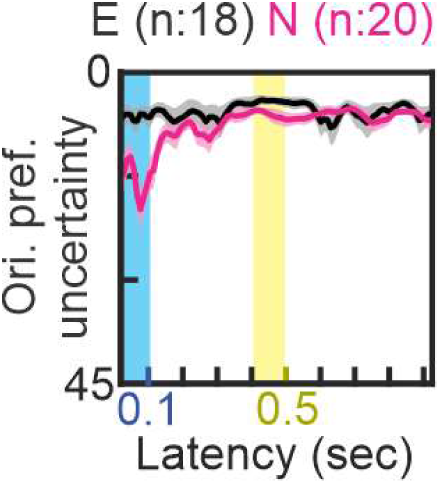
Developmental changes in the temporal consistency of layer 2/3 (L2/3) cells orientation preference is not explained by poor accuracy in preference estimation due to trial-to-trial variability. The plot shows the uncertainty in the estimation of L2/3 sub-threshold response orientation preference as a function of latency following response onset. Uncertainty was computed as the mean error in estimated orientation preference across 100 random re-samplings (with replacement) of single-trial data. Note that the y-axis is inversed to match the format of Figure 4F and responses with higher preference uncertainty, such as low-latency responses in naïve animals, are below responses with lower uncertainty. Uncertainty in the first 100 ms is significantly lower than orientation preference mismatch to the steady response (> 0.5sec, Fig. 4F) for data in experienced (p = 0.003, paired t-test) and naïve animals (p < 0.001).

## Main References

1. Hubel, D.H., and Wiesel, T.N. (1962). Receptive fields, binocular interaction and functional architecture in the cat’s visual cortex. J. Physiol. 160, 106–154.2.

2. Hubel, D.H., and Wiesel, T.N. (1968). Receptive fields and functional architecture of monkey striate cortex. J. Physiol. 195, 215–243.

3. Blasdel, G.G., and Salama, G. (1986). Voltage-sensitive dyes reveal a modular organization in monkey striate cortex. Nature 321, 579. 10.1038/321579a0.

4. Bonhoeffer, T., and Grinvald, A. (1991). Iso-orientation domains in cat visual cortex are arranged in pinwheel-like patterns. Nature 353, 429. 10.1038/353429a0.

5. Malach, R., Amir, Y., Harel, M., and Grinvald, A. (1993). Relationship between intrinsic connections and functional architecture revealed by optical imaging and in vivo targeted biocytin injections in primate striate cortex. Proc. Natl. Acad. Sci. 90, 10469–10473. 10.1073/pnas.90.22.10469.

6. Weliky, M., and Katz, L.C. (1994). Functional mapping of horizontal connections in developing ferret visual cortex: experiments and modeling. J. Neurosci. Off. J. Soc. Neurosci. 14, 7291–7305.

7. Bosking, W.H., Zhang, Y., Schofield, B., and Fitzpatrick, D. (1997). Orientation Selectivity and the Arrangement of Horizontal Connections in Tree Shrew Striate Cortex. J. Neurosci. 17, 2112– 2127. 10.1523/JNEUROSCI.17-06-02112.1997.

8. Douglas, R.J., Koch, C., Mahowald, M., Martin, K.A., and Suarez, H.H. (1995). Recurrent excitation in neocortical circuits. Science 269, 981–985. 10.1126/science.7638624.

9. Li, Y., Ibrahim, L.A., Liu, B., Zhang, L.I., and Tao, H.W. (2013). Linear Transformation of Thalamocortical input by Intracortical Excitation. Nat. Neurosci. 16, 1324–1330. 10.1038/nn.3494.

10. Lien, A.D., and Scanziani, M. (2013). Tuned Thalamic Excitation is Amplified by Visual Cortical Circuits. Nat. Neurosci. 16, 1315–1323. 10.1038/nn.3488.

11. Ko, H., Cossell, L., Baragli, C., Antolik, J., Clopath, C., Hofer, S.B., and Mrsic-Flogel, T.D. (2013). The emergence of functional microcircuits in visual cortex. Nature 496, 96–100. 10.1038/nature12015.

12. Oldenburg, I.A., Hendricks, W.D., Handy, G., Shamardani, K., Bounds, H.A., Doiron, B., and Adesnik, H. (2024). The logic of recurrent circuits in the primary visual cortex. Nat. Neurosci. 27, 137–147. 10.1038/s41593-023-01510-5.

13. Jin, J., Wang, Y., Swadlow, H.A., and Alonso, J.M. (2011). Population receptive fields of ON and OFF thalamic inputs to an orientation column in visual cortex. Nat. Neurosci. 14, 232–238. 10.1038/nn.2729.

14. Chapman, B., Zahs, K.R., and Stryker, M.P. (1991). Relation of cortical cell orientation selectivity to alignment of receptive fields of the geniculocortical afferents that arborize within a single orientation column in ferret visual cortex. J. Neurosci. Off. J. Soc. Neurosci. 11, 1347–1358.

15. Smith, G.B., Hein, B., Whitney, D.E., Fitzpatrick, D., and Kaschube, M. (2018). Distributed network interactions and their emergence in developing neocortex. Nat. Neurosci. 21, 1600– 1608. 10.1038/s41593-018-0247-5.

16. Mulholland, H.N., Kaschube, M., and Smith, G.B. (2024). Self-organization of modular activity in immature cortical networks. Nat. Commun. 15, 4145. 10.1038/s41467-024-48341-x.

17. Feldheim, D.A., and O’Leary, D.D.M. (2010). Visual map development: bidirectional signaling, bifunctional guidance molecules, and competition. Cold Spring Harb. Perspect. Biol. 2, a001768. 10.1101/cshperspect.a001768.

18. Goodhill, G.J. (2016). Can Molecular Gradients Wire the Brain? Trends Neurosci. 39, 202–211. 10.1016/j.tins.2016.01.009.

19. Huberman, A.D., Feller, M.B., and Chapman, B. (2008). Mechanisms Underlying Development of Visual Maps and Receptive Fields. Annu. Rev. Neurosci. 31, 479. 10.1146/annurev.neuro.31.060407.125533.

20. Ackman, J.B., and Crair, M.C. (2014). Role of emergent neural activity in visual map development. Curr. Opin. Neurobiol. 24, 166–175. 10.1016/j.conb.2013.11.011.

21. Chang, J.T., Whitney, D., and Fitzpatrick, D. (2020). Experience-Dependent Reorganization Drives Development of a Binocularly Unified Cortical Representation of Orientation. Neuron 107, 338–350.e5. 10.1016/j.neuron.2020.04.022.

22. Chapman, B., Stryker, M.P., and Bonhoeffer, T. (1996). Development of Orientation Preference Maps in Ferret Primary Visual Cortex. J. Neurosci. 16, 6443–6453. 10.1523/JNEUROSCI.16-20-06443.1996.

23. Trägenap, S., Whitney, D.E., Fitzpatrick, D., and Kaschube, M. (2023). The nature-nurture transform underlying the emergence of reliable cortical representations. Preprint at bioRxiv, 10.1101/2022.11.14.516507 https://doi.org/10.1101/2022.11.14.516507. “The developmental emergence of reliable cortical representations”, Nature Neuroscience, in press.

24. Sellers, K.K., Bennett, D.V., Hutt, A., Williams, J.H., and Fröhlich, F. (2015). Awake vs. anesthetized: layer-specific sensory processing in visual cortex and functional connectivity between cortical areas. J. Neurophysiol. 113, 3798–3815. 10.1152/jn.00923.2014.

25. Schaefer, M.K., Hechavarría, J.C., and Kössl, M. (2015). Quantification of mid and late evoked sinks in laminar current source density profiles of columns in the primary auditory cortex. Front. Neural Circuits 9, 52. 10.3389/fncir.2015.00052.

26. Mitzdorf, U. (1985). Current source-density method and application in cat cerebral cortex: investigation of evoked potentials and EEG phenomena. Physiol. Rev. 65, 37–100. 10.1152/physrev.1985.65.1.37.

27. Chapman, B., and Stryker, M.P. (1993). Development of orientation selectivity in ferret visual cortex and effects of deprivation. J. Neurosci. 13, 5251–5262.

28. Abbott, L.F. (1994). Decoding neuronal firing and modelling neural networks. Q. Rev. Biophys. 27, 291–331. 10.1017/s0033583500003024.

29. Ben-Yishai, R., Bar-Or, R.L., and Sompolinsky, H. (1995). Theory of orientation tuning in visual cortex. Proc. Natl. Acad. Sci. U. S. A. 92, 3844–3848.

30. Chadwick, A., Khan, A.G., Poort, J., Blot, A., Hofer, S.B., Mrsic-Flogel, T.D., and Sahani, M. (2023). Learning shapes cortical dynamics to enhance integration of relevant sensory input. Neuron 111, 106–120.e10. 10.1016/j.neuron.2022.10.001.

31. Murphy, B.K., and Miller, K.D. (2009). Balanced Amplification: A New Mechanism of Selective Amplification of Neural Activity Patterns. Neuron 61, 635–648. 10.1016/j.neuron.2009.02.005.

32. Joglekar, M.R., Mejias, J.F., Yang, G.R., and Wang, X.-J. (2018). Inter-areal Balanced Amplification Enhances Signal Propagation in a Large-Scale Circuit Model of the Primate Cortex. Neuron 98, 222–234.e8. 10.1016/j.neuron.2018.02.031.

33. Pattadkal, J.J., Zemelman, B.V., Fiete, I., and Priebe, N.J. (2024). Primate neocortex performs balanced sensory amplification. Neuron 112, 687–688. 10.1016/j.neuron.2024.01.025.

34. Powell, N.J., Hein, B., Kong, D., Elpelt, J., Mulholland, H.N., Kaschube, M., and Smith, G.B. (2024). Common modular architecture across diverse cortical areas in early development. Proc. Natl. Acad. Sci. U. S. A. 121, e2313743121. 10.1073/pnas.2313743121.

35. Averbeck, B.B., Latham, P.E., and Pouget, A. (2006). Neural correlations, population coding and computation. Nat. Rev. Neurosci. 7, 358–366. 10.1038/nrn1888.

36. Shadlen, M.N., and Newsome, W.T. (1998). The Variable Discharge of Cortical Neurons: Implications for Connectivity, Computation, and Information Coding. J. Neurosci. 18, 3870– 3896. 10.1523/JNEUROSCI.18-10-03870.1998.

37. Zohary, E., Shadlen, M.N., and Newsome, W.T. (1994). Correlated neuronal discharge rate and its implications for psychophysical performance. Nature 370, 140–143. 10.1038/370140a0.

38. Lin, I.-C., Okun, M., Carandini, M., and Harris, K.D. (2015). The Nature of Shared Cortical Variability. Neuron 87, 644–656. 10.1016/j.neuron.2015.06.035.

39. Reinhold, K., Lien, A.D., and Scanziani, M. (2015). Distinct recurrent versus afferent dynamics in cortical visual processing. Nat. Neurosci. 18, 1789–1797. 10.1038/nn.4153.

40. Miller, K.D. (2016). Canonical computations of cerebral cortex. Curr. Opin. Neurobiol. 37, 75–84. 10.1016/j.conb.2016.01.008.

41. Bi, G., and Poo, M. (2001). Synaptic modification by correlated activity: Hebb’s postulate revisited. Annu. Rev. Neurosci. 24, 139–166. 10.1146/annurev.neuro.24.1.139.

42. Clopath, C., Büsing, L., Vasilaki, E., and Gerstner, W. (2010). Connectivity reflects coding: a model of voltage-based STDP with homeostasis. Nat. Neurosci. 13, 344–352. 10.1038/nn.2479.

43. Espinosa, J.S., and Stryker, M.P. (2012). Development and Plasticity of the Primary Visual Cortex. Neuron 75, 230–249. 10.1016/j.neuron.2012.06.009.

44. Sengpiel, F., Stawinski, P., and Bonhoeffer, T. (1999). Influence of experience on orientation maps in cat visual cortex. Nat. Neurosci. 2, 727–732. 10.1038/11192.

45. Tanaka, S., Ribot, J., Imamura, K., and Tani, T. (2006). Orientation-restricted continuous visual exposure induces marked reorganization of orientation maps in early life. NeuroImage 30, 462–477. 10.1016/j.neuroimage.2005.09.056.

46. Blakemore, C., and Cooper, G.F. (1970). Development of the Brain depends on the Visual Environment. Nature 228, 477–478. 10.1038/228477a0.

47. Hughes, N.J., Hunt, J.J., Cloherty, S.L., Ibbotson, M.R., Sengpiel, F., and Goodhill, G.J. (2014). Stripe-rearing changes multiple aspects of the structure of primary visual cortex. NeuroImage 95, 305–319. 10.1016/j.neuroimage.2014.03.031.

48. Paik, S.-B., and Ringach, D.L. (2011). Retinal origin of orientation maps in visual cortex. Nat. Neurosci. 14, 919–925. 10.1038/nn.2824.

49. Ringach, D.L. (2004). Haphazard wiring of simple receptive fields and orientation columns in visual cortex. J. Neurophysiol. 92, 468–476. 10.1152/jn.01202.2003.

50. Schottdorf, M., Keil, W., Coppola, D., White, L.E., and Wolf, F. (2015). Random Wiring, Ganglion Cell Mosaics, and the Functional Architecture of the Visual Cortex. PLOS Comput. Biol. 11, e1004602. 10.1371/journal.pcbi.1004602.

51. Schottdorf, M., Eglen, S.J., Wolf, F., and Keil, W. (2014). Can Retinal Ganglion Cell Dipoles Seed Iso-Orientation Domains in the Visual Cortex? PLOS ONE 9, e86139. 10.1371/journal.pone.0086139.

52. Soodak, R.E. (1987). The retinal ganglion cell mosaic defines orientation columns in striate cortex. Proc. Natl. Acad. Sci. 84, 3936–3940. 10.1073/pnas.84.11.3936.

## Methods references

1. Chang, J.T., Whitney, D., and Fitzpatrick, D. (2020). Experience-Dependent Reorganization Drives Development of a Binocularly Unified Cortical Representation of Orientation. Neuron 107, 338–350.e5. 10.1016/j.neuron.2020.04.022.

2. Scholl, B., Wilson, D.E., Jaepel, J., and Fitzpatrick, D. (2019). Functional Logic of Layer 2/3 Inhibitory Connectivity in the Ferret Visual Cortex. Neuron 104, 451–457.e3. 10.1016/j.neuron.2019.08.004.

3. Chen, T.-W., Wardill, T.J., Sun, Y., Pulver, S.R., Renninger, S.L., Baohan, A., Schreiter, E.R., Kerr, R.A., Orger, M.B., Jayaraman, V., et al. (2013). Ultra-sensitive fluorescent proteins for imaging neuronal activity. Nature 499, 295–300. 10.1038/nature12354.

4. Dana, H., Mohar, B., Sun, Y., Narayan, S., Gordus, A., Hasseman, J.P., Tsegaye, G., Holt, G.T., Hu, A., Walpita, D., et al. (2016). Sensitive red protein calcium indicators for imaging neural activity. eLife 5. 10.7554/eLife.12727.

5. Wilson, D.E., Smith, G.B., Jacob, A.L., Walker, T., Dimidschstein, J., Fishell, G., and Fitzpatrick, D. (2017). GABAergic Neurons in Ferret Visual Cortex Participate in Functionally Specific Networks. Neuron 93, 1058–1065.e4. 10.1016/j.neuron.2017.02.035.

6. Smith, G.B., Hein, B., Whitney, D.E., Fitzpatrick, D., and Kaschube, M. (2018). Distributed network interactions and their emergence in developing neocortex. Nat. Neurosci. 21, 1600– 1608. 10.1038/s41593-018-0247-5.

7. Yang, L., Lee, K., Villagracia, J., and Masmanidis, S.C. (2020). Open source silicon microprobes for high throughput neural recording. J. Neural Eng. 17, 016036. 10.1088/1741-2552/ab581a.

8. Wilson, D.E., Scholl, B., and Fitzpatrick, D. (2018). Differential tuning of excitation and inhibition shapes direction selectivity in ferret visual cortex. Nature 560, 97. 10.1038/s41586-018-0354-1.

9. Brainard, D.H. (1997). The Psychophysics Toolbox. Spat. Vis. 10, 433–436.

10. Lempel, A.A., and Nielsen, K.J. (2019). Ferrets as a Model for Higher-Level Visual Motion Processing. Curr. Biol. 29, 179–191.e5. 10.1016/j.cub.2018.11.017.

11. Lempel, A.A., and Nielsen, K.J. (2021). Development of visual motion integration involves coordination of multiple cortical stages. eLife 10, e59798. 10.7554/eLife.59798.

12. Pettersen, K.H., Devor, A., Ulbert, I., Dale, A.M., and Einevoll, G.T. (2006). Current-source density estimation based on inversion of electrostatic forward solution: effects of finite extent of neuronal activity and conductivity discontinuities. J. Neurosci. Methods 154, 116–133. 10.1016/j.jneumeth.2005.12.005.

13. Sellers, K.K., Bennett, D.V., Hutt, A., Williams, J.H., and Fröhlich, F. (2015). Awake vs. anesthetized: layer-specific sensory processing in visual cortex and functional connectivity between cortical areas. J. Neurophysiol. 113, 3798–3815. 10.1152/jn.00923.2014.

14. Schaefer, M.K., Hechavarría, J.C., and Kössl, M. (2015). Quantification of mid and late evoked sinks in laminar current source density profiles of columns in the primary auditory cortex. Front. Neural Circuits 9, 52. 10.3389/fncir.2015.00052.

15. Mitzdorf, U. (1985). Current source-density method and application in cat cerebral cortex: investigation of evoked potentials and EEG phenomena. Physiol. Rev. 65, 37–100. 10.1152/physrev.1985.65.1.37.

16. Powell, N.J., Hein, B., Kong, D., Elpelt, J., Mulholland, H.N., Kaschube, M., and Smith, G.B. (2024). Common modular architecture across diverse cortical areas in early development. Proc. Natl. Acad. Sci. U. S. A. 121, e2313743121. 10.1073/pnas.2313743121.

17. Kaschube, M., Schnabel, M., Löwel, S., Coppola, D.M., White, L.E., and Wolf, F. (2010). Universality in the Evolution of Orientation Columns in the Visual Cortex. Science 330, 1113– 1116. 10.1126/science.1194869.

18. Mulholland, H.N., Kaschube, M., and Smith, G.B. (2024). Self-organization of modular activity in immature cortical networks. Nat. Commun. 15, 4145. 10.1038/s41467-024-48341-x.

19. Wilson, H.R., and Cowan, J.D. (1972). Excitatory and inhibitory interactions in localized populations of model neurons. Biophys. J. 12, 1–24. 10.1016/S0006-3495(72)86068-5.

20. Wilson, H.R., and Cowan, J.D. (1973). A mathematical theory of the functional dynamics of cortical and thalamic nervous tissue. Kybernetik 13, 55–80. 10.1007/BF00288786.

21. Dahmen, D., Layer, M., Deutz, L., Dąbrowska, P.A., Voges, N., von Papen, M., Brochier, T., Riehle, A., Diesmann, M., Grün, S., et al. (2022). Global organization of neuronal activity only requires unstructured local connectivity. eLife 11, e68422. 10.7554/eLife.68422.

22. Trägenap, S., Whitney, D.E., Fitzpatrick, D., and Kaschube, M. (2023). The nature-nurture transform underlying the emergence of reliable cortical representations. Preprint at bioRxiv, 10.1101/2022.11.14.516507 https://doi.org/10.1101/2022.11.14.516507. “The developmental emergence of reliable cortical representations”, Nature Neuroscience, in press.

23. Abbott, L.F., Rajan, K., and Sompolinsky, H. (2011). Interactions between Intrinsic and Stimulus-Evoked Activity in Recurrent Neural Networks. In The Dynamic Brain: An Exploration of Neuronal Variability and Its Functional Significance, P. Ding Mingzhou and P. Glanzman Dennis, eds. (Oxford University Press), p. 0. 10.1093/acprof:oso/9780195393798.003.0004.

